# An interwoven network of transcription factors, with divergent influences from FoxP3, underlies Treg diversity

**DOI:** 10.1101/2023.05.18.541358

**Authors:** Kaitavjeet Chowdhary, Juliette Léon, Deepshika Ramanan, Diane Mathis, Christophe Benoist

**Affiliations:** Department of Immunology, Harvard Medical School, Boston, MA, USA; INSERM UMR 1163, University of Paris, Imagine Institute, Paris, France

## Abstract

FoxP3+CD4+ regulatory T cells (Tregs), essential for immunologic and organismal homeostasis, have diverse functions and corresponding gene expression programs. How the many controlling transcription factors (TFs) organize to determine Treg identity and diversity remains unclear. We combined single-cell chromatin accessibility profiling, machine learning, and high-density natural genetic variation, validated with TF knockout, CRISPR-editing, and binding data, to define the Treg regulatory network. Distal enhancers proved driven by imbricated multi-TF inputs, employing strategies different from promoter regions. Topic modelling resolved a framework of chromatin programs shaped by distinct TF motifs. This framework anchored surprisingly heterogenous responses to IL2. It identified an unrecognized role for the Smarcc1 remodeler. FoxP3 impacted only some segments of this framework, either activating or repressing programs, amplifying a core Treg identity defined independently. Its absence in Treg-like cells unleashed cytokine expression, but not Th de-differentiation. This work provides a unifying scaffold to understand and manipulate Treg states.

## INTRODUCTION

Cell identities are defined by characteristic gene expression programs. Each program is controlled by the action of specific transcription factors (TFs), which bind and regulate target *cis*-regulatory elements to effect changes in gene expression^1^. TFs act combinatorially, by binding to common regulatory elements, assembling into complexes, or organizing into transcriptional networks. In some systems, so-called “master TFs” sit at the apex of such regulatory networks^2–4^: their expression is both necessary and sufficient to initiate cell-type-specific programs, by launching feedback and feed-forward loops or antagonizing factors that promote alternative fates^1, 5^. Other TFs are required not for cell-type specification but rather for responses to environmental signals^6^. While many studies of how TFs regulate cell identity have focused on transitions between differentiated cell types, how diversification is achieved within a single cell type has received less attention.

Regulatory T cells (Treg) are a subset of CD4+ T lymphocytes that act as dominant controllers of immunologic and organismal homeostasis. Humans and mice with dysfunctional Tregs due to loss-of-function mutations in Foxp3, the Treg lineage-defining TF, develop early-onset, uncontrolled autoimmunity and lymphoproliferation^7^. Tregs have diverse functions, distributed across varied phenotypic poles^8^. For example, distinct Treg programs marked by T-bet+CXCR3+ or IRF4+ phenotypes preferentially restrain Th1 or Th2 inflammation, respectively^9, 10^. Non-lymphoid tissues harbor unique Treg populations^11^, which enforce tolerance to commensal microbes^12^, facilitate tissue regeneration^13^, or control extra-immunologic consequences of inflammation^14^. Treg phenotypic specialization is undergirded by characteristic molecular programs^9, 10, 15–20^. This specialization is often considered in terms of a “one TF-one state” model^7^, in which the expression of single context-specific TFs (e.g. T-bet, PPARγ, RORγ, cMAF, BATF), along with that of FoxP3, mediates differentiation of each Treg subpopulation^9, 19, 21– 25^, although more combinatorial models have been considered as well ^26, 27^. However, how the many TFs expressed in Tregs are systematically organized to determine Treg identity and diversity remains unclear.

Although Foxp3 expression defines Treg identity, its mechanism of action has not been resolved. FoxP3 does not act as a pioneer factor, instead opportunistically binding to regions opened earlier in development^28, 29^. Unlike traditional lineage-defining master regulators, FoxP3 is neither fully necessary nor sufficient to establish Treg identity: Treg-like cells (“Treg wannabes”) can develop in the absence of FoxP3, and FoxP3-independent and -dependent modules characterize Treg-specific gene expression signatures^30–35^. FoxP3 interacts with a large array of transcriptional regulators, chromatin remodelers, and TFs to program Treg identity^36–39^. However, the active structure of FoxP3, and even its recognition motif(s) in DNA are in question^40, 41^, and there is unsettled debate as to whether FoxP3 acts directly as an activator^31, 42–44^, a repressor^36, 37, 45, 46^, or both, depending on its interacting cofactors^39^, or indirectly by tuning the expression of other TFs^47^. An integrative view of how FoxP3 affects TF control across diverse Treg states has so far been elusive.

Here, we integrate several orthogonal strategies to systematically connect TFs to their target Treg programs (Fig 1A). To study the most proximal effects of TF action without confounders from transcriptional bursting or stability, we focused on the role of TFs in modulating chromatin accessibility. Using single-cell chromatin-accessibility profiling, we found that diverse Treg programs were shaped not by individual master TFs, but rather by imbricated multi-TF inputs. Combining machine learning approaches, natural genetic variation, and Treg-specific TF knockouts (KOs), we parsed this combinatorial complexity to resolve the organization of the Treg genetic regulatory network (GRN). FoxP3 had profound and varied effects on this network, with differential influences on distinct chromatin programs. Collectively, these results offer a holistic and clarifying perspective on how one cell type coordinates combinations of TFs and *cis*-regulatory elements to achieve phenotypic diversification.

**Figure 1:**
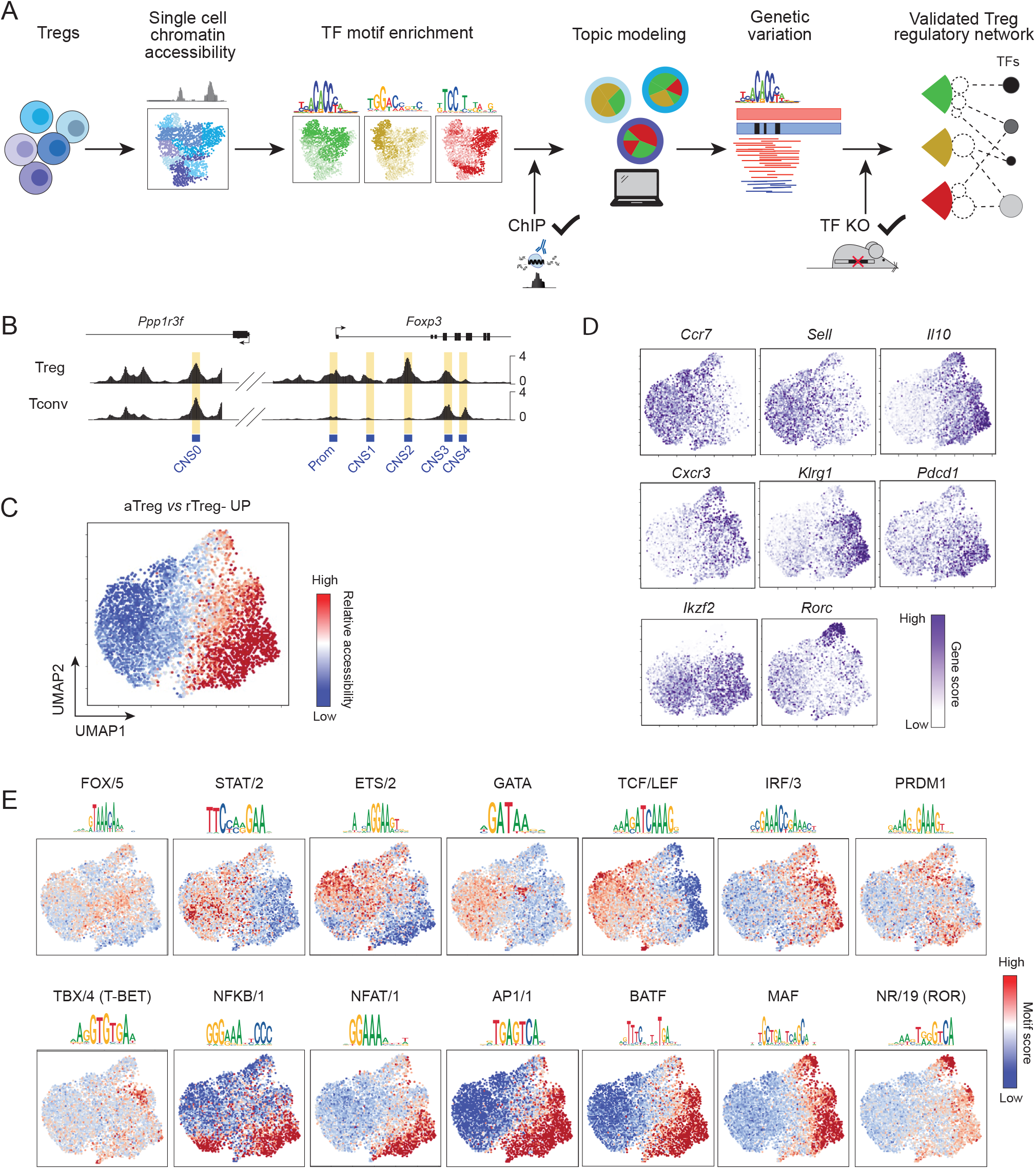
Single-cell ATAC-seq reveals imbricated transcription factor activities across diverse Treg cell states. A) Experimental Overview. Single cell chromatin accessibility profiling was used to link TF activities to diverse Treg chromatin states. Continuous Treg cell states were first annotated using TF motif enrichment. Topic modeling was used to learn groups of co-varying OCRs that formed discrete regulatory modules underlying observed cell states. *Cis*-regulatory variation in B6/Cast F1 hybrid scATAC-seq data enabled identification of causal regulators of Treg chromatin programs. The resulting Treg regulatory network was validated using TF binding (ChIP-seq, CUT&RUN) and knockout datasets. B) Aggregated accessibility profiles of Treg and Tconv single cells at the *Foxp3* locus from scATAC-seq data of splenic Tregs generated from a *Foxp3*^IRES-GFP^ reporter mouse; highlights indicate conserved non-coding sequence (CNS) loci previously described to control *Foxp3* expression. C) Relative accessibility (chromVAR scores) across Treg single cells of OCRs increased in accessibility in aTreg vs rTreg populations (FoldChange>2 in data from ref from^47^) visualized on UMAP of splenic Treg scATAC-seq data. D) Gene scores, chromatin-based proxies for gene expression, visualized for select genes on Treg UMAP from (C). E) Relative accessibility (chromVAR motif scores) of OCRs containing indicated TF motifs; motifs averaged within ‘archetypes’^51^ to reduce redundancy. Only motifs whose corresponding TF(s) are expressed in Treg cells are shown. Motif logos are representative of TFs from each archetype.

## RESULTS

### Imbricated transcription factor activities underlie Treg diversity

To gain a broad view of how accessibility of regulatory regions varies across diverse Treg subpopulations, we generated single-cell Assay for Transposase-Accessible Chromatin using sequencing (scATAC-seq) profiles from splenic Treg (TCRβ+CD4+GFP+) and T conventional (Tconv; TCRβ+CD4+GFP-) cells sorted from a male *Foxp3*^IRES-GFP^ reporter mouse (Fig S1A)^48^. Data were of high quality (median 3x10^4^ fragments per cell, Table S1) and after filtering, we retained profiles for 5,810 Treg and 1,654 Tconv cells (Fig S1B-E). Aggregation of reads from Treg and Tconv single cells recapitulated known patterns of cell-type specific chromatin accessibility at the *Foxp3* locus (Fig 1B). To enable comparisons, we mapped reads to a common peak set consisting of open chromatin regions (OCRs) from a pan-immune cell atlas^29^ supplemented with new peaks identified in this dataset (Table S2).

Within the Treg pool, scATAC-seq profiles captured splenic Treg heterogeneity at multiple levels. Tregs separated broadly by activation status in a 2D uniform manifold approximation and projection (UMAP) visualization, as indicated by relative accessibility of OCR signatures that distinguish activated (aTreg) and resting (rTreg) populations (Fig 1C, Table S3)^47^. To more granularly annotate Treg diversity, we computed ‘gene scores,’ chromatin-based proxies for gene expression^49^. Confirming the chromatin signatures, rTreg and aTreg populations were reciprocally marked by high gene scores for *Ccr7* and *Sell* versus *Il10* (Fig 1D). Importantly, aTregs could be further delineated by gene scores for Treg functional molecules such as *Cxcr3*, *Klrg1*, and *Pdcd1*, each representing previously defined markers of distinct Treg poles^9, 18, 19, 50^. We also noted a small but distinct cluster of cells with high *Rorc* (encoding RORγ) gene scores, corresponding to a Treg subset that dominates in the colon, but is also present at low levels in the spleen^24, 25^. Aggregated accessibility tracks per cell state confirmed the robust differential signals at these loci (Fig S2).

With this landscape of Treg heterogeneity in our scATAC-seq data, we returned to our driving question, how TF activity relates to these Treg states. We examined the relative accessibility per cell of OCRs that contain known TF motifs, grouped into “archetypes”^51^ to reduce redundancy (Fig 1E, S1F). Activation-related motifs had the greatest variability across single cells, cleanly partitioning the Treg pool (Fig S1G). For OCRs with AP-1, NF-AT, or NF-κB motifs, accessibility was highest in aTregs, as expected since these are generically activation-related TFs. However, closer examination revealed imbricated arrangements: each motif was preferentially active in a slightly different region of the aTreg space, and each region included interlaced accessibility of multiple motifs. This perspective synthesized disparate results about individual TFs and their relevance to Treg physiology. For instance, although both c-Maf and RORγ have been reported to be active within the same population, Treg-specific c-Maf knockouts have broader phenotypes^52^. Accordingly, preferential accessibility of the NR/19 motif (corresponding to RORγ) was restricted to only a portion of the large swath of aTregs in which the MAF motif was preferentially accessible. Several studies have suggested BATF to be a main driver of tissue-Treg programs, but its ablation in Tregs affects only a fraction of tissue-Treg-related OCRs^19, 53^. Here, relative accessibility of the BATF motif did not stand out from several other motifs with similar patterns. The activity of OCRs containing TCF/LEF was also curtailed in aTregs, consistent with the dampened expression of TCF-1 and LEF-1, but not with the notion that TCF-1 would be a general FoxP3-controlled mediator of the Treg chromatin program across Treg states^47^. A “one TF-one state” model of Treg diversification would predict that individual TF motifs would be confined to discrete, mutually exclusive cell states. Instead, we found that imbricated and overlapping combinations of multiple factors constitute the diversity of Treg programs.

### OCR usage in Treg single cells

How did this phenotypic variance at the cell level arise from the activity of individual OCRs? To understand the organization of regulatory element activity, we computed similarities in OCR usage across Treg and Tconv single cells, using cosine similarity between outputs from latent semantic indexing (LSI) of OCRs, and visualized them in a 2D UMAP projection (Fig 2A). OCR activity patterns did not resolve into clearly defined classes, instead forming graded continua. Most OCRs overlapping or near transcription start sites (TSS) separated strikingly from those mapping further away (‘distal’ OCRs, mostly in enhancers) (Fig 2B, Fig S3A). Consistent with prior observations^29, 54–56^, TSS OCRs had the least variable accessibility (Fig 2C), confirming that distal OCRs contributed most to Treg diversity. Cell- and state-specific OCRs grouped together on the OCR UMAP visualization (Fig 2D, Table S3). This analysis therefore demonstrated that individual OCRs had unique and independent activity patterns and were not simply co-induced as discrete regulatory blocks.

**Figure 2:**
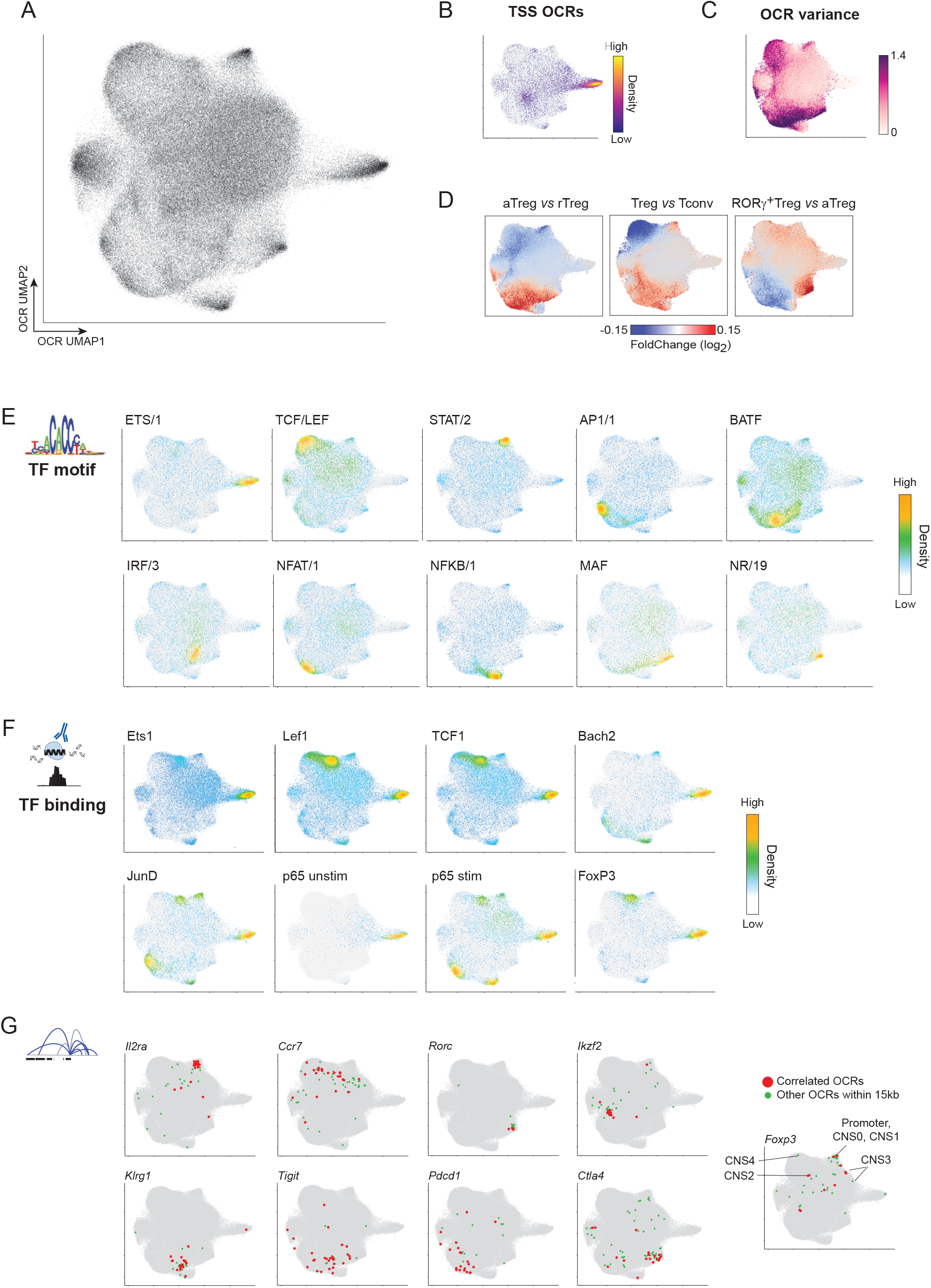
Global usage of open chromatin regions across Treg single cells. A) UMAP visualization of OCR usage across Treg and Tconv single cells from *Foxp3*^IRES-GFP^ spleen scATAC-seq data in Fig 1. B) OCR UMAP from (A) with OCRs overlapping annotated TSS highlighted; color indicates density of TSS OCRs. C) OCR UMAP from (A) colored by variance in accessibility across Tregs. D) UMAP from (A) colored by log_2 F_old Change between indicated populations from Treg scATAC-seq data in Figure 1. E) Annotation of OCRs overlapping indicated TF motif archetypes on OCR UMAP from (A); color indicates density of motif-containing OCRs. F) Annotation of OCRs overlapping indicated TF binding sites on OCR UMAP from (A); color indicates density of bound OCRs. G) Annotation on OCR UMAP from (A) of OCRs with accessibility correlated (FigR^71^ p < 0.05) with expression of indicated genes (in red) or other OCRs within 15 kb of gene TSS not meeting this correlation significance threshold (in green). Locations of *Foxp3* conserved non-coding sequences (CNS) and promoter loci are indicated.

Next, we asked how TFs interfaced with this organization of Treg OCRs. As noted in other contexts^51, 57^, individual OCRs contained motifs for several TFs, with a median of 8 motifs per OCR (Fig S3B). To connect TF binding to OCR activity, we positioned their motifs onto the OCR UMAP (Fig 2E). Motifs were not present at random, but demarcated distinct (STAT, NF-κB, and NR/19 (RORγ)) yet overlapping (NF-AT, AP-1, BATF) accessibility patterns in the OCR space. We also observed variegated configurations when projecting the binding sites of several TFs, deduced from ChIP-seq and CUT&RUN experiments in Tregs^28, 47, 58–61^ (Fig 2F, Table S4). TFs bound preferentially but not exclusively to distinct groups of state-specific OCRs, with different TFs occupying rTreg-(Lef1, TCF-1) or aTreg-(Bach2, JunD) biased loci. Binding distributions were dynamic: in unstimulated Tregs, the NF-κB component p65 bound primarily to TSS regions, but in stimulated cells shifted to occupy diverse distal regions (Fig 2F), including a cluster of OCRs enriched for NF-κB motifs (Fig 2F). FoxP3-binding had a narrow footprint on the OCR landscape (Fig 2F), with greatest enrichment among TSS and a group of rTreg-preferential distal OCRs, a concentration among clusters of STAT and NF-κB motif-containing OCRs, and diffuse minor representation among aTreg-specific loci. Thus, mirroring results at the cell level, each group of OCRs was occupied by overlapping combinations of TFs.

How did organization of OCR usage relate to gene expression? Most genes are controlled by multiple *cis*-regulatory elements^62–64^. This multiplicity confers molecular and evolutionary robustness, but also enables the expression of one gene in different differentiated cell types^65–67^. Our OCR UMAP allowed us to ask, within a single cell type, how OCRs linked to the same gene varied in their patterns of accessibility, and hence shared regulatory drivers. Several techniques have been proposed to connect OCRs to their target genes^29, 64, 68–71^. We formed OCR-gene links by using covariation of OCR accessibility with gene expression from paired splenic Treg scATAC and single cell transcriptomic (scRNA) datasets (FDR < 0.05)^71^, also annotating OCRs within 15kb of each TSS although not meeting correlation criteria (Fig 2G, Fig S4, Table S5). Visualization of gene-linked OCRs on the OCR UMAP revealed divergent regulatory strategies. While some genes had OCRs with homogeneous activity patterns (e.g., *Rorc, Klrg1*), others had more varied distributions (e.g., *Ccr7*) or even multiple sub-patterns (e.g., *Tigit, Pdcd1*, *Ctla4*), with disparate accessibility profiles among loci previously shown to control *Foxp3* expression^72^. Thus, genes vary in how flexibly their associated OCRs are used across Treg states.

### Topic modeling learns Treg chromatin programs

If individual TFs were insufficient to parse the imbricated organization of the Treg GRN, could one instead group patterns of OCR usage into co-regulated modules, and then determine the TFs that drove them? Each cell state within the Treg continuum could be conceptualized as the integration of multiple discrete regulatory programs. To learn such programs, we employed a machine learning approach, probabilistic topic modeling, derived from text mining, to categorize co-varying OCRs into “topics”^73^. Topic modeling, which has been applied to single cell genomic data^74, 75^, is well-suited for sparse and incomplete data and, unlike conventional partition clustering approaches, allows for the attribution of more than one program to each OCR, thus better accommodating the continuous structure of the Treg OCR space. In practice, we modified a previous method for topic modeling of scATACseq data^74^ for robustness by using an ensemble approach to create a consensus set of reproducible topics. The optimal solution parsed 17 such topics (Computational Note 1). Each topic learned by this strategy captured a different pattern of coordinated accessibility across Treg single cells. OCRs could belong to multiple topics, each topic being strongly represented in 1.5 to 3.9x10^4^ OCRs (Computational Note 1, Table S6). OCRs from each topic occupied distinct poles of the OCR UMAP space (Fig 3A) and had diverse patterns of accessibility across Treg single cells (Fig 3B). Some topics, which included an overrepresentation of TSS OCRs (e.g., 6 and 8), were broadly accessible, while others captured highly specific gradations in accessibility. For example, Topics 3, 10, and 14 learned groupings of OCRs modulated in different facets of the aTreg pool, separate from Topic 9, which included OCRs active in RORγ+ cells.

**Figure 3:**
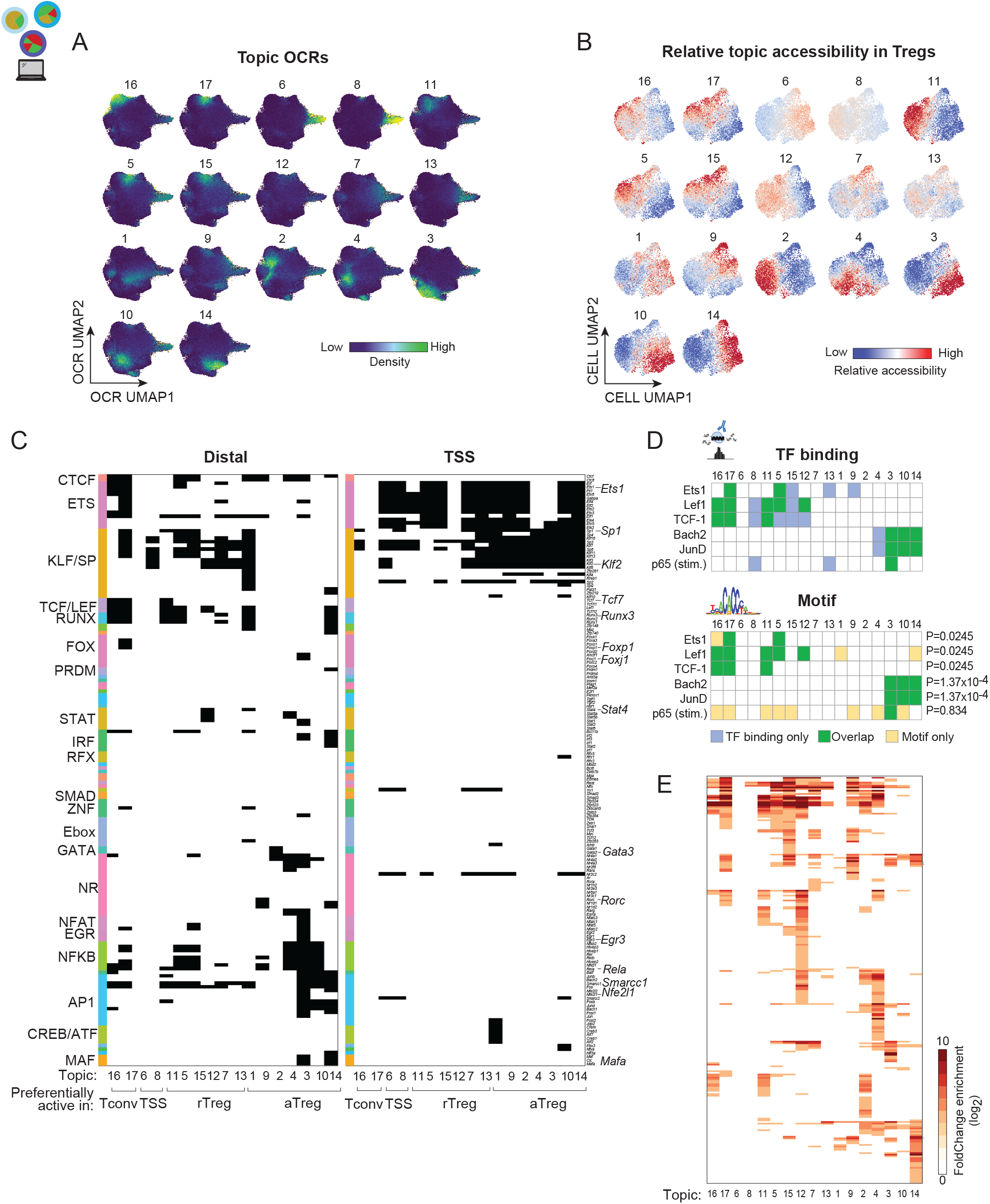
Topic modeling learns coordinated accessibility programs across Treg single cells. A) Annotation of OCRs assigned to each topic on OCR UMAP from Fig 2; color indicates density of OCRs. B) Relative accessibility (chromVAR scores) of OCRs from each topic across Treg single cells, visualized on UMAP of Treg scATAC-seq data from Fig 1. C) Enrichment of TF motifs within distal or TSS OCRs from each topic; black indicates enrichment FDR < 1x10^-10^. Motifs are organized by TF family (row annotation). D) Overlap between enrichment of TF-binding data (top) or corresponding motifs (bottom). Green indicates concordant enrichments in both motif and TF-binding analyses, whereas blue indicates enrichment only in TF-binding data and yellow enrichment only using motif data. P value indicates significance of overlapping enrichments between motif and binding data (binomial test). E) Gene Ontology gene sets significantly enriched among regulatory regions in each topic (using GREAT ^76^ analysis). Heatmap indicates fold change of enrichment relative to background. Full table of enrichments and pathway names is provided in Table S7.

Thus, topic modeling provided a quantitative approach to summarize patterns of Treg OCR accessibility and a tractable entry point to relate Treg chromatin programs to TF activity. To nominate candidate regulators of topic OCR accessibility, we computed the enrichment of TF motifs within each topic, yielding very different assignments for TSS vs distal OCRs (Fig 3C, Table S7). Distal OCR enrichments highlighted state-specific TF connections. While topics more accessible in aTregs were enriched in motifs for AP-1, NF-κB, and nuclear receptor factors, Tconv- and rTreg-preferential topics instead included KLF/SP, ETS, and TCF/LEF motifs (consistent with ^47^). Some enrichments were highly specific: Gata3 only in Topic 2, RORγ only in Topic 9. Importantly, no topic was defined by enrichment of any one TF or TF family, indicating that this modular description of the Treg regulatory program itself required the combinatorial activity of different TFs. Consistent with their less variable accessibility, TSS OCRs were enriched in motifs represented in most topics, including ETS and KLF/SP family members.

To validate these links between TFs and topics, we made use of independent experimental TF-binding data. Using TFs for which high quality ChIP-seq or CUT&RUN profiles in Tregs were available, we compared the predicted topic-specific motif enrichments with the distribution of biochemical TF binding sites (Fig 3D). This overlap was highly significant for almost all TFs examined. Thus, enrichment of TF motifs within topics delineated the structure by which combinatorial TF inputs connected to diverse Treg epigenomic programs.

We used GREAT^76^ to connect OCRs to genes, and identify functional pathways enriched in the regulatory regions defined by each topic (Fig 3E,Table S7). We will refrain from cherry-picking the results, but the over-arching conclusion was that the Topics did not merely bring up scattered functional annotations (which they might have) but different biological programs – for instance, it is not as if all aTreg-associated Topics flagged the same activation-related gene sets. Topic 14 had distinctive associations related to regulation of other immunocytes.

Overall, these topics now provide a framework that can be re-applied to compare the state of the Treg GRN in different datasets and experiments. This is illustrated for several experiments below, and we have implemented a web application (https://cbdm.connect.hms.harvard.edu/Topic_Plotting/) that allows any investigator to upload a single-cell dataset and receive its decomposition in terms of these Topics.

### Topics across tissues

Tregs traffic to and take residence in nonlymphoid tissues (tissue Tregs), where they elaborate specialized regulatory programs^11^. As the multi-step acquisition of tissue-Treg modules begins in lymphoid organs^16, 17, 19, 20, 50, 77^, it was important to know whether topics learned in spleen Tregs would capture tissue-Treg programs. To start, we looked for the distribution of a set of OCRs that we had previously reported as being induced in tissue Tregs (colon, visceral adipose tissue, muscle) relative to splenic Tregs^16^ (Table S3). These pan-tissue-Treg OCRs were enriched specifically within Topics 3 and 14 (Fig 4A), with 40% of this OCR set overlapping with these two topics. To further this comparison, we generated new scATAC-seq profiles of Tregs isolated from spleen and colonic lamina propria from the same *Foxp3*^IRES-GFP^ reporter mice (Fig S5A-B), multiplexed by hashtagging in the same run^78^. We computed the variance explained in this new dataset by the topics determined from our initial data. Highlighting the robustness of these topics, there was excellent concordance (Pearson r=0.99) between the relative variance explained across the new and previous spleen Treg scATAC datasets (Fig 4B). Spleen and colon Tregs differed in this regard, however. Topics 3 and 14 explained more variance (Fig 4C) and had increased accessibility within colonic Treg cells (Fig 4D), confirming the earlier enrichment results. Accessibility of Topics 3 and 14 was uniformly increased across colon Treg cells (Fig 4E). Several other topics, mostly biased towards rTreg (e.g., Topics 11, 12, 5), were decreased in accessibility and relative variance explained (Fig 4D, Fig S5D). Topics also captured more specific patterns of accessibility: for example, Topics 9 and 10 were preferentially active in RORγ+ and Helios+ Tregs, respectively (Fig 4F, Fig S5C-D, Table S3). Thus, topics were robust in their explanatory power, and captured tissue-specific specializations, confirming the notion that tissue-Treg programs represent amplifications of patterns already present in Tregs from lymphoid tissues^16, 17, 19, 20, 50, 77^. Together with Fig. 3C, which indicates that Topics 3 and 14 were not driven by any one TF but by an ensemble, these results indicate that tissue-Treg chromatin programs are contained within the topic framework and result from combinatorial opening.

**Figure 4:**
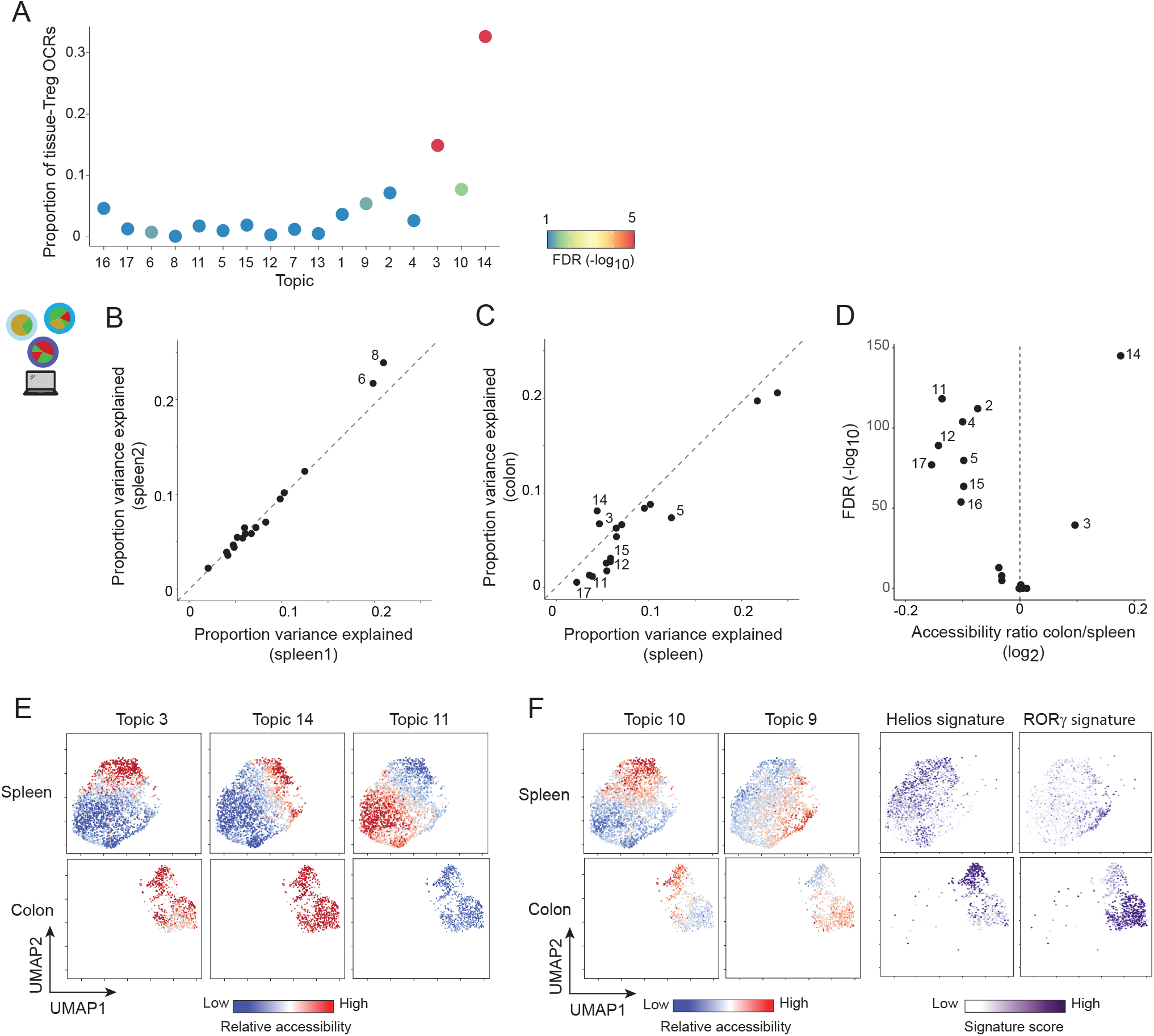
Topics across tissues. A) Proportion of pan-tissue Treg OCRs (from ^16^) overlapping each Treg topic; color indicates significance of enrichment (permutation test). B) Proportion of variance in accessibility explained by each topic in spleen scATAC data from Fig 1 (spleen 1) or in independently generated *Foxp3*^IRES-GFP^ spleen Treg scATAC data in Figure 4 (spleen 2). C) Proportion of variance in accessibility explained by each topic in colon or spleen Treg scATAC data (from the same *Foxp3*^IRES-GFP^ reporter mouse). D) Differential accessibility per topic between aggregated colon and spleen Treg scATAC profiles. E) Relative accessibility (chromVAR scores) of OCRs from tissue-(Topics 3, 14) or spleen-(Topic 11) specific topics across spleen and colon Treg single cells visualized on scATAC UMAP, separated by organ. F) Relative accessibility (chromVAR scores) of OCRs from Helios-(Topic 10) or RORγ-(Topic 9) specific topics across spleen and colon Treg single cells visualized on scATAC UMAP, separated by organ. Right panel indicates gene module scores for genes from a Helios vs RORγ specific gene expression signature visualized on the same UMAP.

### Topics and the response to IL2

Having established the Treg chromatin programs at baseline in lymphoid and nonlymphoid tissues at steady state, it was of interest to assess how this network would adapt to acute stimulus. IL2 and the STAT family TFs that it predominantly activates, are canonical controllers of Treg cell differentiation in the thymus, and of the maintenance of FoxP3 expression and Treg numbers in the periphery ^79–82^, in a homeostatic negative feedback loop ^83–86^. There have been detailed studies of IL2’s transcriptional signature in Tregs ^87, 88^.

We generated scATAC-seq profiles of splenic Tregs from mice treated acutely (2 hrs prior to favor direct effects without secondary confounders) with IL2 (Fig. S6A). Underscoring the potent effects of IL2, this short exposure led to a marked shift in Treg chromatin states (Fig 5A). After classifying cells as rTreg and aTregs based on their relative accessibility of OCR signatures from Fig 1 (Fig 5B, S6B), the UMAP visualization (Fig 5A,B, S6C) showed that rTreg populations shifted more than did aTreg cells in response to IL2 (median Local Inverse Simpson’s Index^89^ between treated and untreated 1.13 in rTreg vs 1.60 in aTreg, p<2.2x10^-16^; Fig S6D). Visualization of the closest cells in treated vs untreated pools in high-dimensional OCR space (nearest neighbor in an LSI embedding with IL2 effect removed ^89^) confirmed that all rTregs responded sharply but only a fraction of aTregs did (Fig. 5C). This lesser response of aTregs was also reflected by lower STAT5 phosphorylation after exposure to IL2 in culture (Fig. 5D), and explainable at least in part by a lower presence of the high affinity receptor for IL2, IL2Ra, on the surface of aTregs (Fig. 5E), with lower accessibility of enhancers upstream and within the gene body of the *Il2ra* locus in aTregs (Fig. S6E). Induction of STAT motif-containing OCRs was seen across rTreg and aTreg responses (Fig S6F,G). aTregs had lower levels of STAT target accessibility at baseline and following stimulation (Fig S6G). However, while rTregs homogeneously increased their relative chromatin accessibility of STAT5-containing OCRs in response to IL2, aTregs responded heterogeneously (Fig. 5F). Smiegel et al previously reported a lower responsiveness to IL2 among CD44^hi^CCR7^lo^Tregs ^90^, attributing this low reactivity to an inability of those cells to use CCR7 to home to locales of high IL2 concentration. Since the injected IL2 in our experiments is diffusible and not constrained by local production, the present results indicate an intrinsically low response of aTregs to IL2, relative to rTregs, and suggest that they may be less sensitive to IL2-centered therapeutic interventions.

**Figure 5:**
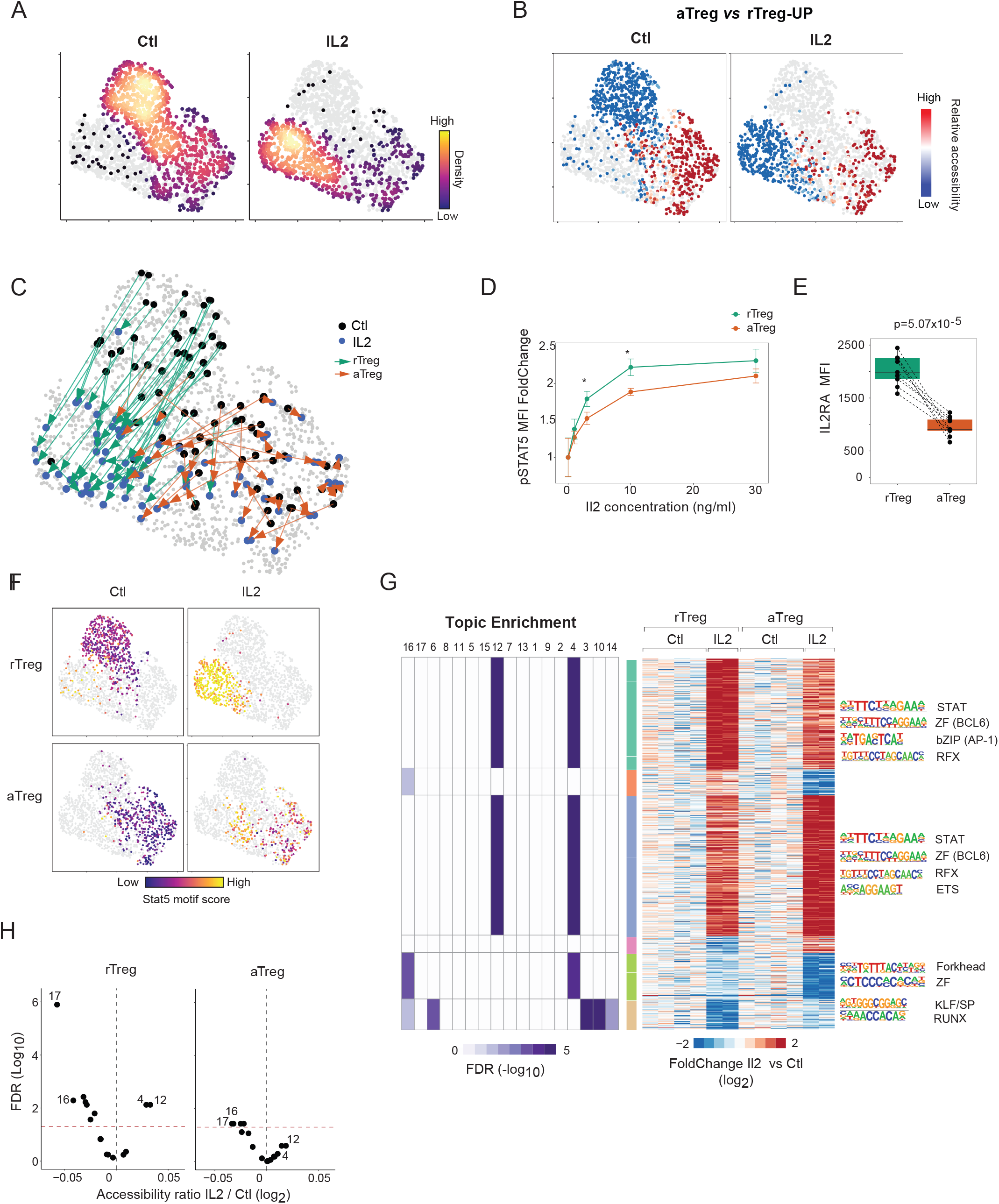
Topics and the response to IL2. A) UMAP of scATAC-seq of splenic Tregs from *Foxp3*^IRES-GFP^ mice treated with 10 μg of recombinant mouse IL2 (IV) or PBS vehicle control for 2 hours, colored by density of cells from each condition. B) Relative accessibility (chromVAR scores) of OCRs increased in accessibility in aTreg vs rTreg populations visualized on UMAP of scATAC-seq data from (A) and split by treatment group. C) Visualization of pairs identifying the closest cells in IL2 -untreated and -treated conditions in high-dimensional OCR space, overlaid onto UMAP from (A), and colored by the cell state (rTreg or aTreg) of the untreated, control cell in each pair. Random samples of 40 pairs of rTreg control and 40 pairs of aTreg control cells shown for visual clarity. D) Fold Change of rTreg or aTreg pSTAT5 mean fluorescence intensity, relative to untreated control, after *ex vivo* stimulation with indicated concentration of recombinant mouse IL2 for 15 minutes. p values from t.test. N=3 mice. E) IL2RA (CD25) mean fluorescence intensity among splenic rTreg or aTreg. p value from paired t.test. N=9 mice. F) Relative accessibility of OCRs containing Stat5 motif (chromVAR scores), visualized on UMAP from A and split by cell state and treatment status. G) Heatmap of OCRs with |log_2F_oldChange|>1.5 in accessibility between treated and untreated conditions in either rTreg or aTreg comparisons (860 OCRs). Displayed values indicate Fold Change (log_2)_ of aggregated accessibility in treated versus untreated cells, matched for cell state (rTreg or aTreg). Each column indicates a different biological replicate. Heatmap to left shows significance of enrichment (log_10(_FDR), permutation test) of Topic OCRs in each OCR cluster (enrichments with FDR<0.05 shown). TF motifs (HOMER ^144^, p value < 10^-5^) enriched in each OCR cluster are indicated on right. Values underlying results provided in Table S8. H) Differential accessibility per topic between aggregated IL2 treated vs untreated spleen Treg scATAC profiles, separated by cell state.

As might be expected, no topic uniquely encompassed the response to IL2, and only a minority of OCRs were affected within each topic. Selecting robust responses (|log2FoldChange|>1.5 in either rTregs or aTregs (860 OCRs), we clustered the different patterns of changes in accessibility (Fig. 5G, Table S8), and determined the match between Topics and these clusters (Fig. 5G, left panel and Table S8). Induced OCRs were specifically enriched for Topics 4 and 12. Topic 12 contains an interferon-responsive component (Fig 3E), and its induction was consistent with recent data indicating that IL2 elicits a significant Interferon-Stimulated Gene response in Tregs ^88^, most likely because IL2 signals via STAT1 as well as STAT5. Motif enrichment showed these OCRs to be enriched for STAT5 and BCL6 motifs. (Fig 5G, right panel and Table S8). Repressed OCRs, on the other hand, were enriched for Topic 16 regions across all cells, confirmed by an integrated analysis that also revealed a weaker but more general effect on Topic 17 OCRs (Fig. 5H). Topics 16 and 17 are chromatin programs preferentially active in Tconv, which suggests that IL2 bolsters Treg identity by suppressing Tconv-specific features, a notion of interest given IL2’s role in supporting Treg differentiation in the thymus ^85^. OCRs with greater reduction in accessibility among rTregs were enriched in aTreg-specific topics (Topics 3, 10, 14), and KLF/SP or RUNX motifs, whereas those with greater reduction in aTregs were enriched in Forkhead or ZF motifs. Thus, chromatin topics captured the specific response to IL2, described differences across cell states, and highlighted connections to different inductive and repressive mechanisms of IL2 action.

### Natural genetic variation causally parses state- and OCR-specific TF effects

To bolster and validate these patterns derived from machine learning, we exploited naturally occurring genetic variation as an orthogonal approach to functionally establish causal relationships. Wild-derived Cast/Eij (Cast) mice differ from reference C57BL/6 (B6) mice by approximately 20 million variants^91^. As elegantly established in previous work^47, 92–96^, B6xCast F1 offspring can be used to causally link sequence variation to chromatin features: because the two genomes are present within the same cell, controlling for any changes in *trans* effects (i.e., TF expression), allelic skews in chromatin accessibility can be causally attributed to *cis-*regulatory alterations (i.e., disrupted TF motifs), the equivalent of a genome-wide mutagenesis experiment.

We generated scATAC profiles of Tregs sorted from B6xCast F1 mice (Fig S7A-B; n=5,980 Tregs). We avoided known biases in reference-specific read mapping by adapting a published pipeline^97^ to align reads to a common coordinate system, assigning informative reads to their allele of origin (Fig S7C). We adapted analytical frameworks established in previous studies (“mean diff” calculations^47, 95^) to link changes in TF motifs with corresponding shifts in allele-specific chromatin accessibility (“allelic motif effect (AME)”), with an algorithmic modification to apply to the topic modeling context (Computational Note 2). For each topic, we identified the cells with accessibility of topic OCRs containing each candidate motif and computed the F1 AME for motifs in each topic only in these relevant cells (Fig 6A). Thus, in contrast to bulk analyses of F1 data that identify *average* effects of TFs genome-wide and in all cells, the resulting topic-specific AME quantified the causal contribution of TF motifs to each individual topic, only in the relevant cells in which the topic is active.

**Figure 6:**
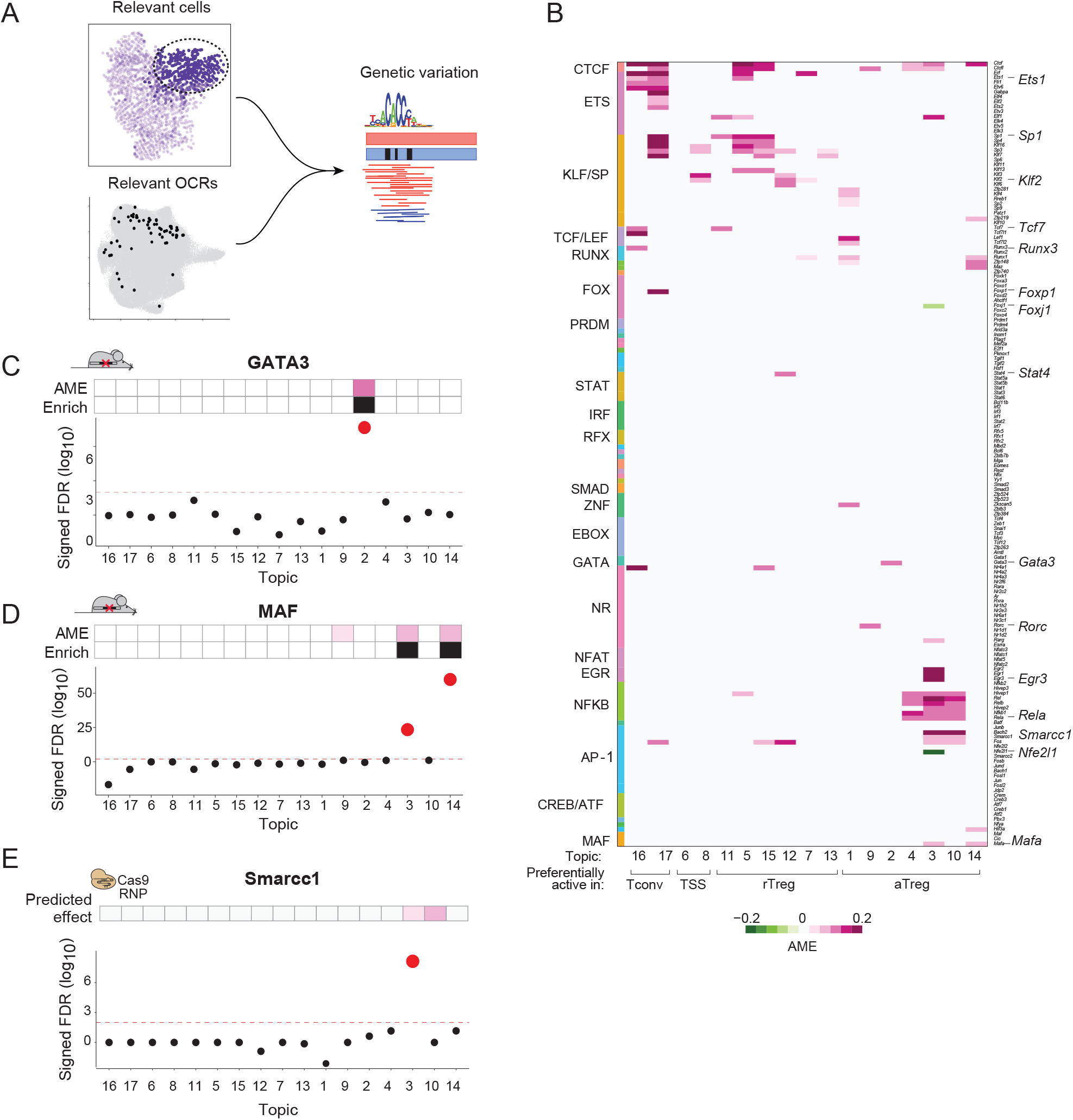
Genetic variation identifies causal regulators of Treg chromatin programs. A) Overview of Topic AME calculation: to quantify the contribution of motifs to the accessibility of each topic, AME was calculated in reads aggregated from cells with high accessibility of topic OCRs containing each candidate motif in spleen Treg scATAC-seq profiles from B6/Cast F1 hybrid mice. B) Topic-specific AME (FDR<0.10) of motifs in each Treg topic. Heatmap in same order as in Figure 3 and shows overlap between motif enrichment and significant AME scores. Positive AMEs (pink) indicate positive effect on chromatin accessibility and negative AMEs (green) indicate negative effect on chromatin accessibility. Motifs are ordered by TF family. C) Enrichment (signed log_10(_FDR), permutation test) in each topic of Gata3-dependent OCRs (GATA-motif containing OCRs decreased in accessibility > two-fold in Treg-specific Gata3 KO vs WT). Panel above indicates predicted topic effect based on topic AME (top bar) or GATA family motif enrichment (bottom bar). D) Enrichment (signed log_10(_FDR), permutation test) in each topic of c-Maf-dependent OCRs (MAF-motif containing OCRs decreased in accessibility > 2-fold in Treg-specific c-Maf KO vs WT). Panel above indicates predicted topic effect based on MAF family topic AME (top bar) or motif enrichment (bottom bar). E) Enrichment (signed log_10(_FDR), permutation test) in each topic of Smarcc1-dependent OCRs (OCRs with loss of accessibility at p<0.05 in *Foxp3*^IRES-GFP^ Tregs electroporated with CRISPR/Cas9 ribonucleoprotein complexes carrying *Smarcc1*-targeting versus control gRNAs and transferred for 1 week into Treg-depleted *Foxp3*^DTR^ hosts). Panel above indicates predicted topic effect based on combination of topic AME and motif enrichment from 6B.

The results provided an unprecedented view of regulators of Treg chromatin programs, both in breadth and context-specificity. To identify the strongest modulators of accessibility, we highlighted effects detected in both motif enrichment and AME analyses (Fig 6B, Table S9). However, motifs with significant AMEs without corresponding enrichment may still reflect causal regulatory function (all AMEs in Fig S7D). The vast majority of AMEs corresponded to positive contributions of TF motifs to chromatin accessibility, very few motifs (e.g., Foxj1, Nfe2l1) having repressive effects, with members of the same TF family at times having opposing actions. While ETS, KLF/SP, and TCF/LEF motif families had detectable AMEs in several topics, their genetic effects overlapped with motif enrichments primarily in rTreg- and Tconv-biased topics. AMEs sharpened the scope of AP-1 and NF-κB effects, narrowing their multi-topic enrichment to a cluster of AMEs specific to aTreg-preferential Topics 3, 4, and 10. AMEs also identified restricted effects: for example, in accordance with its accessibility in the small group of RORγ+ cells, Topic 9 had a significant AME for the RORγ motif. Thus, integrating genetic effects with topic motif enrichment pierced through the combinatorial imbrication of the Treg regulatory network to refine links between TFs and Treg chromatin programs.

We validated this Treg TF network by testing with orthogonal *trans*-regulatory perturbations. First, we evaluated the impact of Treg-specific Gata3 ablation by generating scATAC profiles of Tregs from *Foxp3-cre*×*Gata3^fl/fl^* mice and *Foxp3-cre*×*Gata3^+/+^*littermates. Gata3-dependent OCRs defined from these data (GATA motif-containing OCRs with a greater than two-fold decrease in accessibility in the knockout; Table S3) were enriched only in Topic 2 (FDR < 0.01), the topic predicted to be under Gata3 control based on both AME and motif enrichment (Fig 6C). Secondly, we used published ATAC-seq data from Tregs sufficient or deficient in c-Maf^98^. In our model, MAF family motifs had significant AMEs in Topics 9, 3, and 14 but enrichment only in Topics 3 and 14. Accordingly, c-Maf-dependent OCRs (Table S3) from the knockout analysis were enriched only in Topics 3 and 14 (FDR < 0.01; Fig 6D). Thus, TF elimination in Tregs validated and matched predictions of TF-dependent control from our Treg TF network.

Thirdly, we tested the importance of a regulatory factor predicted by the model but not previously associated with Treg biology. Smarcc1 (BAF155) is a core subunit of all mammalian SWI/SNF chromatin remodeling complexes ^99, 100^. Some SWI/SNF components have been associated with changes during T cell activation ^96, 101^ or regulation of *Foxp3* expression ^102^, but Smarcc1 has not been previously implicated in control of state-specific Treg chromatin. Our network predicted a specific effect of Smarcc1 in aTreg-skewed topics. To test its relevance, we delivered CRISPR-Cas9 ribonucleoprotein complexes ^103^ carrying *Smarcc1*-targeting or control gRNAs to CD4+ T cells isolated from *Foxp3*^IRES-GFP^ mice, parked *in vivo* the edited cells in Treg-depleted *Foxp3*^DTR^ hosts, and sorted GFP+ Tregs 1 week later for bulk ATAC-seq analysis. OCRs which lost accessibility in Smarcc1 KO cells (p<0.05, Table S3) were specifically enriched (FDR<0.01) for Topic 3 OCRs, as predicted by our network (Fig 6E). Topic 10 was not represented, potentially reflecting redundancy between Smarcc1 and Smarcc2 paralogs ^99, 104^. Thus, based on the network predictions, we experimentally validated an unrecognized role for Smarcc1 in selectively controlling aTreg-specific chromatin programs.

### FoxP3-independent and -dependent control of the Treg TF network

While the above provided an integrative view of TF control of Treg chromatin programs, the elephant in the room was FoxP3, the Treg lineage-defining TF, which did not appear in our network. This result is consistent with the uncertainties that surround the DNA motif(s) actually recognized by FoxP3^40, 41^ and the notion that FoxP3 is not a pioneer factor that modifies chromatin accessibility^28, 29^. To understand the intrinsic role of FoxP3 in a setting unconfounded by systemic inflammation, we made use of female mice heterozygous for a Foxp3 loss-of-function allele (Foxp3^fs327^–GFP/Foxp3-Thy1.1 mice, “KO” in Fig 7A)^44^. As Foxp3 is encoded on the X-chromosome, due to random X-inactivation, one population of Treg cells expresses wild-type FoxP3 protein (flagged by the Thy1.1 reporter), while another population of Treg-like cells expresses a Foxp3 allele with a full loss-of-function frameshift mutation whose expression is reported by GFP. The presence of functional Thy1.1+ Tregs prevents immune dysregulation, thus providing a well-controlled system for investigating FoxP3-intrinsic effects. Control mice (WT in Fig 7A) are similarly constructed, but with a functional FoxP3 encoded upstream of the GFP reporter.

**Figure 7:**
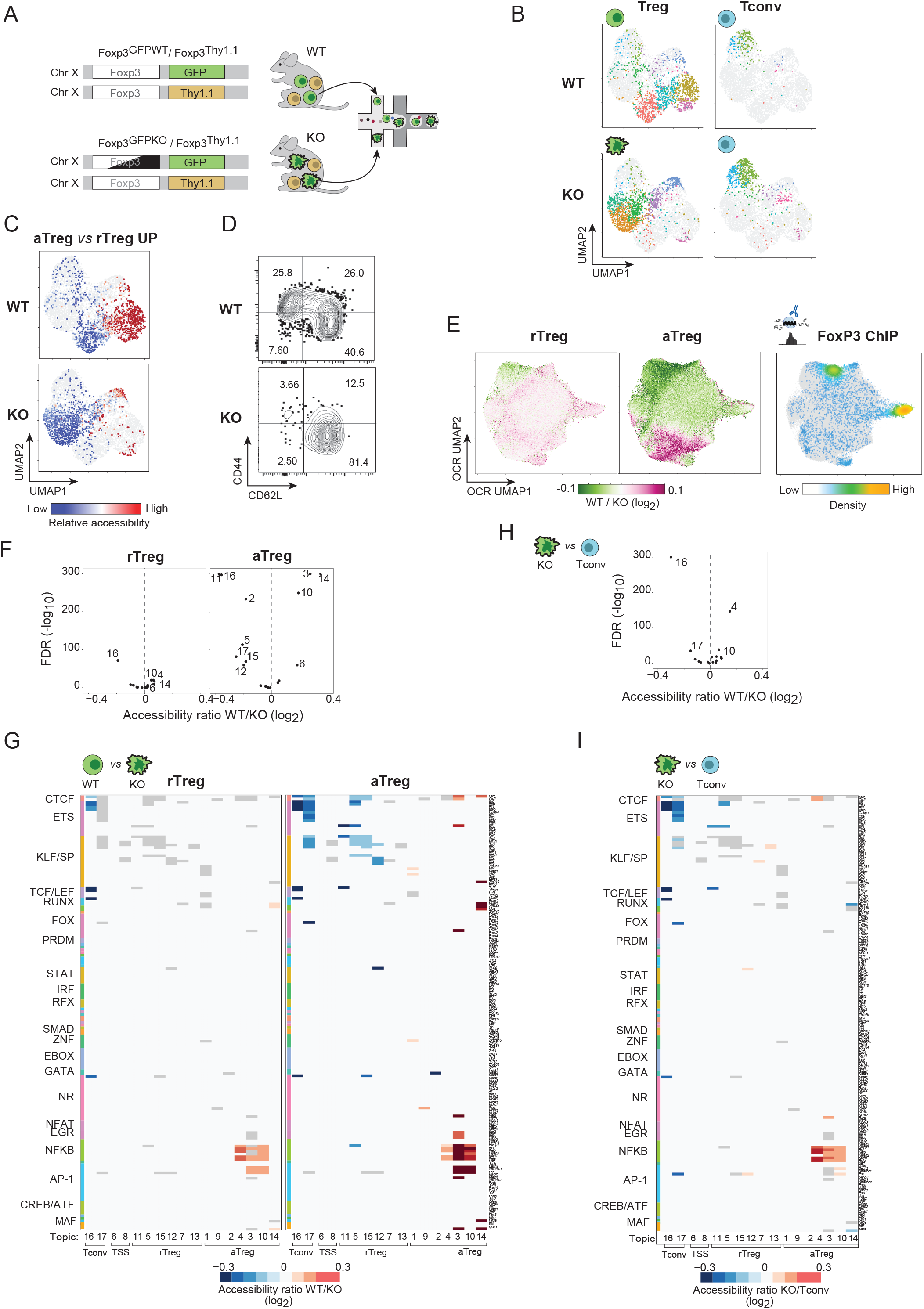
FoxP3 effects on the Treg regulatory network. A) Experimental Scheme. FoxP3-deficient (KO) and -sufficient (WT) GFP+ Tregs were sorted along with GFP-Tconv from Foxp3^fs327^–GFP/Foxp3-Thy1.1 or Foxp3^wt^-GFP/Foxp3-Thy1.1 heterozygous female mice for scATAC-seq. B) UMAP of scATAC of Treg and Tconv from FoxP3 WT or KO heterozygous female mice, separated by genotype and cell type. C) Relative accessibility (chromVAR scores) across WT and KO Treg single cells of OCRs increased in accessibility in aTreg vs rTreg populations, visualized on UMAP of scATAC-seq data from (B). D) Proportion of rTreg and aTreg populations in FoxP3 WT or KO populations by CD44 and CD62L flow cytometry. E) OCR UMAP from Fig 2 colored by log_2 F_oldChange in chromatin accessibility in FoxP3 WT vs KO cells in aTreg or rTreg comparisons. Right panel shows annotation of OCRs overlapping FoxP3 binding sites; color indicates density of FoxP3-bound OCRs. F) Differential accessibility per topic between WT Treg and KO Treg-like cells in rTreg or aTreg comparisons. G) Differential accessibility per motif in each topic (distal OCRs, FDR<0.05) between WT and KO Treg cells in rTreg or aTreg comparisons for motif to topic connections from Figure 6B. Grey indicates significant motif to topic connection from topic AME in Figure 6B but no significant change in accessibility across FoxP3 comparisons. H) Differential accessibility per topic between KO rTreg and Tconv cells. I) Differential accessibility per motif in each topic (distal OCRs, FDR<0.05) between KO rTreg and Tconv cells for motif to topic connections from Figure 6B. Grey indicates significant motif to topic connection from topic AME in Figure 6B but no significant change in accessibility across differential comparisons.

We sorted GFP+ Treg and GFP-Tconv from both WT and KO heterozygous mice for scATAC-seq, hashtagging all cells into the same run by genotype and bins of CD25 expression (Fig 7A-B, Fig S8A-C). UMAP visualization of these data showed profound changes in the chromatin states of FoxP3-deficient GFP+ cells (Fig 7B, Fig S8D). While Tconv from WT and KO mice mostly co-mingled, as expected from the unperturbed environments, FoxP3-deficient Treg-like cells occupied a region of the embedding distinct from that occupied by WT Tregs (Fig 7B). For scale, the Local Inverse Simpson’s Index was significantly lower (p<10^-9^) for FoxP3 KO and WT cells (median=1.21) than for Gata3 KO and WT cells (median = 1.73) (Fig S8E). FoxP3-deficient Treg-like cells also showed diminished accessibility of aTreg-specific OCRs, most KO Tregs being in a resting-like chromatin state (Fig 7C). This inability of FoxP3-deficient Tregs to progress to activated states was confirmed by flow cytometry (Fig 7D, Fig S8F).

To better understand the OCRs driving this shift, we computed differential accessibility between Treg-like cells and WT Tregs, performing comparisons separately in rTreg and aTreg populations to avoid cell composition-driven effects (Table S3). We overlaid the differential accessibility from each comparison onto the OCR UMAP from Fig 2. FoxP3-repressed OCRs (lower accessibility in WT) were concentrated in rTreg- or Tconv-preferential loci, while FoxP3-potentiated OCRs (higher accessibility in WT) were in aTreg-specific regions (Fig 7E). Both at the level of individual OCRs (Fig 7E, S9A) and topics (Fig 7F), FoxP3-dependent changes were more pronounced in comparisons within aTregs than in rTregs, especially for FoxP3-potentiated OCRs. While FoxP3 binding was somewhat more enriched in repressed OCRs, most FoxP3-dependent changes were unrelated to FoxP3 binding (Fig 7E, Fig S9A), supporting previous results^28, 47^. Compared with FoxP3-negative loci, FoxP3-bound OCRs were more widely accessible (Fig S9B) and less variable (Fig S9C) across Treg single cells.

How did FoxP3 influence the Treg GRN? For a comprehensive view, we looked for differential accessibility within motif-topic connections defined above (Fig 7G, Table S9). FoxP3 effects split into two groups: FoxP3 repressed ETS, KLF/SP, TCF/LEF, RUNX, and FOX motif accessibility in Tconv- and rTreg-preferential topics, but boosted accessibility of EGR, NF-κB, AP-1, and MAF motifs in aTreg-biased topics (Fig 7G, Fig S9D). Notably, some motifs with causal effects in several topics (e.g., CTCF, ETS, KLF/SP) were affected in opposite directions in different programs, highlighting that FoxP3 did not influence TFs homogenously genome-wide, and underscoring the power of parsing context-specificity with topic modeling. In short, this analysis suggested that FoxP3 had two roles: (1) repressing Tconv- and rTreg-like programs and (2) promoting activation-related chromatin features.

If FoxP3 does not entirely define Treg identity ^31–35, 105, 106^, what are the “Treg wannabes” that develop in its absence? We compared, in the Topic framework, KO Treg-like cells and Tconv. KO Treg-like cells most strongly induced Topic 4 (controlled by NF-κB, known to be important to Treg identity ^107, 108^) and repressed Tconv-preferential Topics 16 and 17 (Fig 7H). Strikingly, these FoxP3-independent effects (Fig. 7I, S9E) seemed a carbon-copy of the FoxP3-dependent effects observed when comparing WT and KO Tregs (Fig. 7G, Fig. S9F). Together, these results provide an integrated vista of Treg identity and FoxP3 function. Core features of Treg identity are established independently of FoxP3 and subsequently amplified by its expression. FoxP3 is then required for Treg activation, where FoxP3 protects Treg identity and enables the induction of aTreg-specific chromatin programs, which underly Treg suppressive and effector functions.

### FoxP3 deficiency differentially affects Treg subsets in vivo

We noted an overrepresentation of cells with high *Rorc* gene scores and NR/19 motif accessibility among the FoxP3-deficient Treg-like population (Fig 8A, Fig S10A-B), which suggested a gut connection for these cells, since RORγ+ Tregs dominate in the colon^24, 25^. The comparison of RORγ+ Treg proportions in KO Treg-like cells with Tregs from WT littermates showed a modest increase in RORγ+ cells across several organs, but a major shift in the colon, where almost all were RORγ+ (Fig 8B). In contrast, a drop in Helios+ Tregs was observed in the colon (Fig 8B-C), consistent with a drop in *Ikzf2* accessibility in the genomic data Fig S10A-B). These results confirm recent observations from van der Veeken et al^109^.

**Figure 8:**
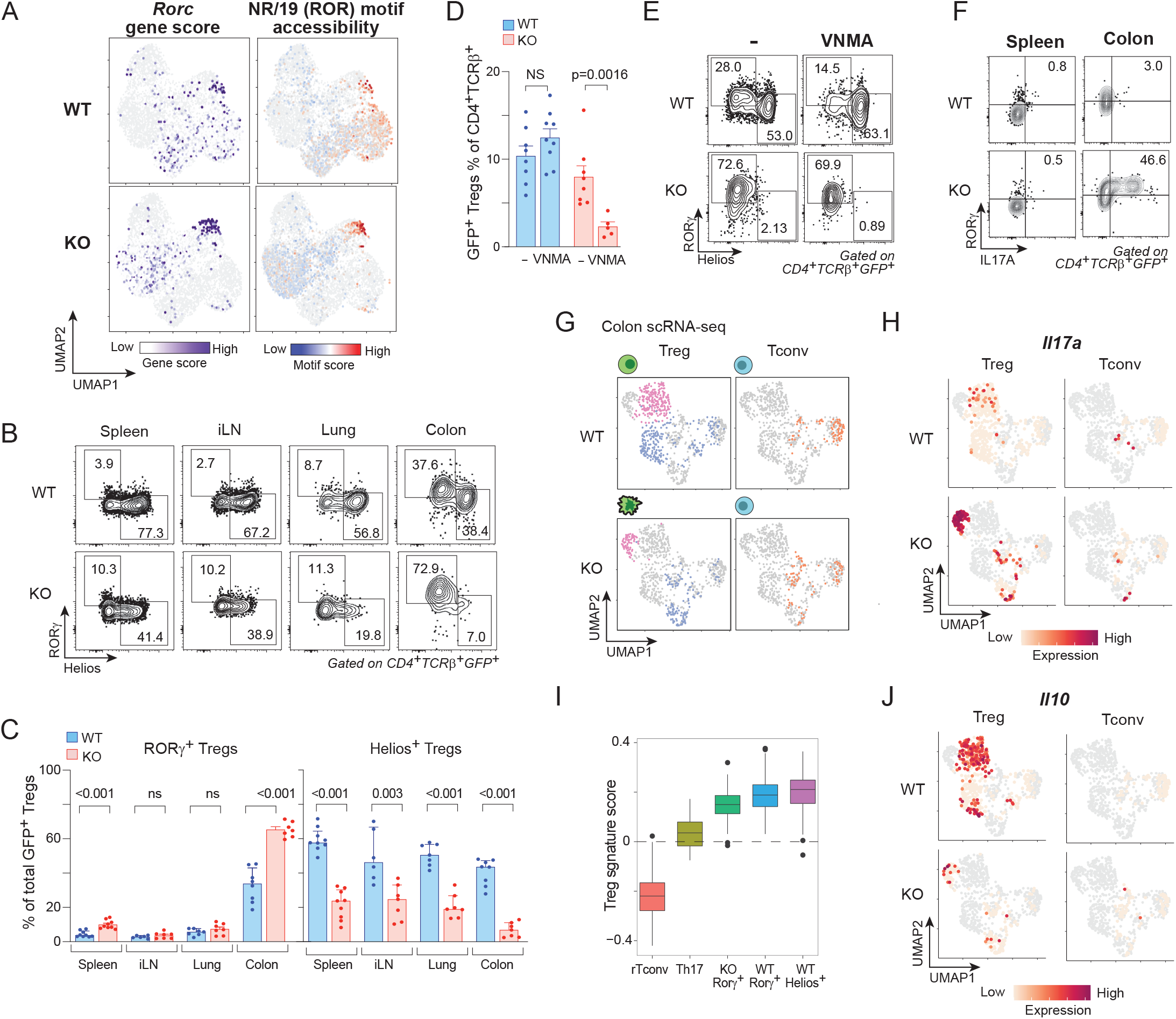
FoxP3-independent RORγ+ Treg-like cells. A) *Rorc* gene scores and NR/19 (contains RORγ) motif accessibility (chromVAR score) in WT and KO Treg populations in scATAC-seq data from Fig 7. B) Proportion of RORγ+ vs Helios+ Tregs among WT or KO Tregs in heterozyous female mice across organs. C) Quantification of (A). D) Proportion of GFP+ Tregs among WT or KO Tregs with or without VNMA antibiotic treatment. E) Proportion of RORγ+ vs Helios+ Tregs among WT or KO colon Tregs before or after VNMA antibiotic treatment. F) Proportion of IL17A+ RORγ+ cells among stimulated FoxP3 KO or WT Tregs from spleen or colon. G) UMAP of scRNA-seq of colonic lamina propria Treg and Tconv from FoxP3 WT or KO heterozygous female mice, separated by genotype and cell type. Pink indicates *Rorc*+ GFP+ WT or KO Tregs, blue indicates all other WT or KO Tregs, and orange indicates Tconv. H) *Il17a* expression overlaid onto UMAP from (G). I) Distribution of gene signature expression scores across indicated populations in colon scRNA-seq data from (G) J) *Il10* expression overlaid onto UMAP from (G).

In normal mice, RORγ+ Tregs are microbiota-dependent^24, 25^, and we asked whether RORγ+ Tregs of KO mice were also regulated by bacterial inputs. After treatment with a broad-spectrum antibiotic cocktail (VNMA, 4 weeks), the fraction of total GFP+ Tregs decreased strongly in the KO colon, but not in WT (Fig. 8D) where the drop in RORγ+ Tregs was balanced by an increase of the Helios+ pool (Fig. 8E). The persisting Treg-like cells in KO mice remained RORγ^+^, and no Helios+ Tregs emerged (Fig. 8E). Thus, confirming and extending the single-cell genomics, this analysis added another layer to the variegated function of FoxP3: RORγ^+^ and Helios+ Treg populations depend differentially on FoxP3.

FoxP3-deficient RORγ+ Tregs in the colon produced IL17A (Fig. 8F), as also recently observed ^109^. This was not the case in the spleen, indicating that the expression of Teff cytokines is not merely unleashed by the absence of FoxP3, but requires an active driver, present in the gut but not in the spleen (Fig 8F). Did these FoxP3-deficient RORγ+ cells maintain their Treg identity, or were they turning into Th17? We performed scRNAseq on sorted GFP+ Treg and GFP-Tconv from colons of WT and KO heterozygous females (Fig 8G). In accordance with the scATACseq, colonic KO Tregs were shifted in the UMAP relative to WT Tregs (Fig 8G, S10C) and had high *Il17a* expression (Fig 8H). However, these KO Tregs clearly remained in the Treg space on the UMAP, far from the *Rorc+Il17a+* Th17 cells in the same dataset (Fig 8G-H, S10C). They retained the ability to express IL10 (Fig. 8J), maintained Treg signature scores closer to WT Treg than to Tconv (Fig 8I), and matched WT Tregs in their *Foxp3* locus accessibility pattern (Fig. S10D). Thus, KO Tregs did not become Th17 cells but maintained several facets of Treg identity while simply de-repressing cytokine production. Notably, another population of *Gata3*+ KO Treg-like cells downregulated *Ikzf2* expression and expressed type 2 cytokine transcripts (*Il4, Il5, Il13*; Fig S10C), indicating parallel de-repression in *Rorc+* and *Gata3+* populations. In summary, while not required for RORγ+ Treg differentiation, FoxP3 was required to repress Teff cytokines induced in the gut environment.

## DISCUSSION

A growing body of work has described a panoply of functions performed by Tregs and characterized their associated molecular programs^11, 110^. Some transcriptional modules that accompany this specialization have been defined, but a systematic view of how TFs are integrated to control these programs has been missing. Here, we combined the informative power of single-cell genomics, machine learning, and high-density genetic variation, with validation by gene knockout and TF-binding datasets, to resolve the architecture of the Treg regulatory network. By avoiding confounders that result from averaging across all cells or enhancers, and by causally linking TF activities to specific chromatin subprograms, the results greatly simplify the nebulous complexities of Treg control and resolve some of the contradictions. We found that the diversity of Treg chromatin states arose from combinatorial, imbricated gradients of TF inputs. Against this background, FoxP3 had a profound impact, with a diversity of influences on different programs.

Treg enhancers were organized into a continuum, indicating the unique activity of each individual regulatory element across single cells (Fig 2). This was somewhat surprising, as one might have expected that enhancers operational within a defined cell-type would tend to fall into a limited number of classes. This continuum was structured, however: OCRs that bound a given TF largely congregated in particular sections of the OCR space. While the sparsity inherent to scATAC-seq may certainly contribute to the apparent continuity, one might also speculate that it represents cell-to-cell fluctuations in genomic activity (i.e. biological noise ^111^) or dynamic responses to environmental signals^6^. Seemingly subtle modulation of T cell enhancers can lead to dramatically divergent immunological effects^112^. The fine-grained diversity in enhancer usage may enable Tregs to flexibly respond to varied and unpredictable immunological perturbations.

Conceptually or experimentally handling such a diversity of enhancer elements is challenging. Individual TFs, with their overlapping binding patterns, provide insufficient resolution. Topic modeling, although an approximation of these continuous activities, delivered a tractable abstraction through which to learn the drivers of major Treg regulatory programs, and single-cell resolution enabled identification of programs active only in rare subpopulations (e.g., RORγ+ spleen Tregs). Many motifs had inconstant effects across topics, indicating that TFs do not contribute homogeneously to chromatin activity across Treg cells. Such variation in TF operation indicated that Treg TF activity was highly context-specific, perhaps mediated by availability or activity of different cofactors. Parsing TF actions by topic should help resolve the ambiguities and contradictions around the function of individual factors, whose specific actions might otherwise have been obscured by averaging across cells or OCRs in population-level chromatin studies. Topics provided a quantitative summary of Treg chromatin state that proved portable across replication studies and conditions. Topic activities were robustly reproducible and, although learned from baseline spleen data, captured programs amplified in non-lymphoid tissues and parsed heterogeneous responses to IL2 stimulation.

Genetic variation analysis enabled interrogation of causal relationships between TF motifs and Treg topics. Notably, each topic consisted of contributions from some motifs which were important for several topics and other motifs which were highly specific in their activity. This may suggest that the constellation of Treg chromatin programs is driven by combinations of a common set of TFs required for Treg identity overlaid by a specific set of TFs that enable Treg diversity, as observed in other contexts^96^. The power of genetic variation to identify unrecognized regulators was highlighted by our validation of a previously unrecognized role for Smarcc1in the control of a specific aTreg chromatin program.

Motif enrichments were markedly different at OCRs in promoter regions vs distal enhancers. TSS OCRs involved a limited set of motifs, with topic diversification that might occur through coupling with distal sites in the same program, via physical enhancer-promoter interaction. Indeed, analysis of FoxP3-mediated enhancer-promoter loops previously showed distinct motif localization to either side of enhancer-promoter loop structures^44^.

The combinatorial organization of TFs that control each program matched findings from several other fields and organismal settings^57, 65, 113^ but challenged the persistent notion specific to the immunology literature that phenotypic specialization is mediated by single master TFs. For example, several reports have proposed that a particular TF critically controls programs relevant to tissue Tregs (e.g., BATF or Blimp-1^19, 53, 114, 115^), or controls discrete components (e.g., T-bet^9^). Tissue Treg programs did prove dominated by a pair of topics (3 and 14), which do include AP-1 and PRDM family motifs among their controlling regulators, but importantly not in a unique manner, several other TFs conspiring to control these tissue Treg topics. Accordingly, although relative accessibility of the BATF or PRDM1 motifs was highest in aTregs, their distributions overlapped with several other motifs (e.g., other AP-1, MAF, or NF-κB factors) with similar patterns. We speculate that, by extension, the framework developed here might explain why factors identified as controlling pan-tissue-Treg chromatin programs affect only a portion of those OCRs^19,20^

What is the mechanism by which FoxP3 controls Treg regulatory programs? Even in a setting unconfounded by inflammation, FoxP3 deficiency had a major impact on chromatin accessibility. While previous studies have debated whether FoxP3 acts as an activator or repressor, the present results highlight that it acts as both. Importantly, FoxP3’s repressive and activating functions affected distinct chromatin topics with different motif enrichments (Fig 6). The OCR membership of each of these topics provides genomic definition to the locations where FoxP3 might assemble into different nuclearly segregated molecular complexes with distinct co-factor composition and opposing functions^39^.

The results also offer an integrated model for the relationship between FoxP3 and Treg identity. In rTregs, FoxP3 amplified a program of Treg identity present in “Treg wannabes”, with a striking superimposition between the pre-existing program and what FoxP3 amplifies, down to the finest topic and motif details. How this concordance is achieved molecularly is an open question. One interpretation is that FoxP3 is uniquely attuned to cooperate with the TF ensemble that supports the Treg-like wannabe status, or that it replaces a similar Forkhead factor that would anchor the Tconv-wannabe differentiation. Which factors specify the pre-FoxP3 Treg program will be an important area of future study, for which these data provides candidates. FoxP3 affected a greater number of OCRs within aTregs, suggesting an accentuated role for FoxP3 following Treg maturation. FoxP3 deficiency led to disappearance of aTreg populations and decreased accessibility of aTreg-specific OCRs, in accordance with^30, 32, 53^. In principle, this loss of aTregs could be because FoxP3 is required for aTreg differentiation^30^, or because of a diversion of aTregs to an alternative fate (including death) in the absence of FoxP3^32, 116^. These widened FoxP3 effects may depend on the availability of new cofactors in aTregs. Notably, the OCRs most strongly potentiated by FoxP3 belonged to tissue-Treg topics (3 and 14), consistent with the notion that FoxP3 is required for VAT Treg specification^17^, but not colon RORy+ Tregs^109^.

The discovery of a population of FoxP3-independent RORγ+ Tregs added another layer to the relationship between FoxP3 and Treg differentiation. While this manuscript was in preparation, van der Veeken et al also reported FoxP3-independent RORγ+ Tregs^109^. We found FoxP3-deficient RORγ+ Treg-like cells to be microbe-dependent, as are normal FoxP3-sufficient counterparts. They produced IL17A but only in the colon, and without diversion to Th17, suggesting that cytokine de-repression in the absence of FoxP3 requires a positive environmental input, possibly via TCR or Wnt/β-catenin signaling, both of which have previously been associated with RORγ+IL17+ phenotypes^117–119^. However, extrinsic inputs cannot be the sole determinants of this program, as an intrinsic bias towards an increased RORγ/Helios ratio remained even in the context of antibiotic treatment. Thus, RORγ may partially compensate for the absent FoxP3 in these cells, ensuring their competitive survival over Helios+ counterparts, but unable to fulfill all of FoxP3’s functions, like repression of cytokine genes upon stimulation.

Taken together, this study provides a model where TF function is conditioned by cell state, where specific combinations of TFs integrate input signals to enable Treg diversification. FoxP3 plays multimodal functions to maintain Treg identity and enable aTreg and consequently tissue-Treg-specific differentiation. In the colon, Tregs amplify pre-existing TF networks to adapt to new environments. Tissue-specific stimulation presents a unique challenge to self-reactive Tregs, where FoxP3 is important for balancing proportions of RORγ+ and Helios+ Tregs and preventing production of inflammatory cytokines. Overall, these results offer a quantitative and clarifying picture of the TF control of Treg diversity and provide a roadmap for its manipulation.

## Supporting information

Computational Notes

## ACKNOWLEDGEMENTS

We thank R. Ramirez, S. Mostafavi, J. Buenrostro, A. Rudensky, J. van der Veeken, and Y. Zhong for helpful discussions; N. Ramirez, C. Gerhardinger at the Harvard Bauer core for scATAC-seq; the HMS Biopolymers Sequencing Facility; M. Sleeper, J. Nelson, D. Ischiu at the HMS Immunology Flow core; K. Hattori for assistance with mouse strains; C. Laplace for assistance with graphics; D. Mallah for assistance with web tool development. This work was funded by grants from the NIH to C.B. and D.M. (AI150686, AI165697, AI125603) and a grant from the JPB Foundation to C.B. and D.M. K.C was supported by NIGMS grants T32GM007753 and T32GM144273 and a Harvard Stem Cell Institute MD/PhD Training Fellowship, J.L. by an IMAGINE MD/PhD grant and an Arthur Sachs scholarship, and D.R. by the Damon Runyon Cancer Research Foundation (DRG 2300-17, National Mah Jongg League Fellow).

## MATERIALS & METHODS

### Mice

*Foxp3*^IRES-GFP^ mice^48^ and *Foxp3^fs327^*^-GFP^ x *Foxp3^tm10^*^.1(Casp9,-Thy1)Ayr^ (or Foxp3^Thy1.1^)^44^ heterozygous female mice were bred on the C57Bl/6J background and maintained in our colony in the specific pathogen–free facility at Harvard Medical School (HMS). Gata3 conditional knockout mice were generated by crossing Cre+ *Foxp3*-*cre* mice^120^ with heterozygous Gata3*^fl/+^* mice^121^. To generate B6/Cast F1 mice, Cast/EiJ males (Jackson Labs, strain #000928) were crossed with C57BL/6J females (Jackson Labs, strain #000664). All experimentation was performed following animal protocols approved by the HMS Institutional Animal Use and Care Committee (protocols IS00000054 and IS00001257). Except when specified, 6-10-week-old male mice were used throughout this study. For heterozygous FoxP3 KO experiments, 5-10-week-old female mice were used. Experiments involving knockout mice always used WT littermate controls for comparisons.

### Isolation, analysis, and sorting of T lymphocytes

#### Spleen and lymph nodes

Immunocytes were released using mechanical disruption followed by filtering and washes in phenol red-free DMEM containing 10mM HEPES (Gibco) and 2% fetal calf serum (FCS). Red blood cells in spleen samples were lysed before filtering using ACK lysis buffer (Gibco, ref A10492-01). In some samples, CD4+ T cells were enriched using negative magnetic selection using the Dynabeads Untouched Mouse CD4 Cells Kit (ThermoFisher Scientific, cat#11415D).

#### Colon

Immunocytes were isolated as previously described^122^. Briefly, intestines were cleaned (Peyer’s patches removed in the case of the small intestine), and treated with RPMI containing 1 mM DTT, 20 mM EDTA and 2% FCS at 37°C for 15 min to remove epithelial cells. They were then minced and dissociated in collagenase solution (1.5 mg/mL collagenase II (Gibco), 0.5mg/mL dispase (Gibco) and 1% FCS in RPMI) with constant stirring at 37°C for 40 min. Single cell suspensions were filtered and washed with RPMI containing 5% FCS.

#### Lungs

Immunocytes were isolated as previously described^123^. Briefly, mice were first perfused with 5mL of ice-cold PBS through the heart’s right ventricle. Tissues were minced and dissociated in collagenase solution (1 mg/mL collagenase IV (Gibco), 150μg/mL DNase I (Sigma) and 1% FCS in DMEM) and incubated in a water bath at 37°C with constant shaking for 30 min. Digested tissues were filtered and washed in 2% FCS. For lungs, red blood cells were lysed using ACK lysis buffer.

#### Flow Cytometry

After Fc blocking and Live/Dead staining (Zombie Fixable Viability Kit, Biolegend), extracellular staining was done in ice cold phenol red-free DMEM containing 2% FBS for 30 min using antibodies against surface markers (see antibodies below). In order to maintain GFP signal, cells were first pre-fixed in 1% formaldehyde for 15 min at room temperature and then fixed overnight at 4°C using 100 μL of Fix/Perm buffer (eBioscience FoxP3/TF Staining Buffer Set, cat# 00-5523-00). After membrane permeabilization using 1X permeabilization buffer (eBioscience FoxP3/TF Staining Buffer Set, cat# 00-5523-00) for 5 min, intracellular staining was performed for 2 hours at room temperature (see antibodies below). Data was recorded on a FACSymphonyTM (BD Biosciences) or on an Aurora (Cytek Biosciences) flow cytometer and analyzed using FlowJo 10 software.

#### Sorting

Cells were sorted using BD MoFlo Astrios EQ, FACSAria-561, or FACSAria-594 machines. For experiments with *Foxp3*^IRES-GFP^ mice, Tregs were sorted as DAPI-Dump (CD19, Cd11c, Cd8)-CD4+ TCRb+ GFP+ and Tconv (where applicable) as DAPI-Dump (CD19, Cd11c, Cd8)-CD4+ TCRb+ GFP-. For experiments using mice without Foxp3 reporters, Tregs were sorted as DAPI-Dump (CD19, CD11c, CD8)-CD4+ TCRb+ CD25hi and Tconv (where applicable) as DAPI-Dump (CD19, CD11c, CD8)-CD4+ TCRb+ CD25lo. For FoxP3 heterozygous female experiments, KO or WT Treg cells were sorted as DAPI-Dump (CD19, CD8)-CD4+ TCRb+ Thy1.1-GFP+, separating cells by bins of CD25 expression (2 bins for KO and 3 bins for WT cells see Fig S6A-C).

#### Antibodies

For sorting or flow analyses of lymphoid and non-lymphoid tissues, the following antibodies were used (BioLegend): CD4-APC (clone GK1.5) or CD4-BV605 (clone RM4-5), TCRb-PECy7/BUV737/AF700 (clone H57-597), CD25-PE/APC (clone PC-61), CD19-FITC/PB (clone 6D5), CD8a-FITC/PB (clone 53-6.7), CD11b-FITC (clone M1/70), CD44-BV510 (clone IM7), CD62L-BV785 (clone MEL-14), CD45.2-AF700 (clone 104), Thy1.1-PE/PECy7/APC (clone OX-7), RORγ PE/APC/BV785 (clone AFKJS-9), Helios-PB (clone 22F6), IL17A-PE/APC (clone TC11-18H10.1).

### Antibiotic Treatment

For antibiotic treatment, vancomycin (0.5 g/L, VWR Life Science), metronidazole (1 g/L, Sigma), neomycin (1 g/L, Fisher), and ampicillin (1g/L, Sigma) were dissolved in drinking water with 0 calorie sweetener and given to mice for 4 weeks, with replenishment of fresh antibiotics every 7-10 days.

### Cytokine Staining

For *ex-vivo* cytokine staining, single-cell suspensions of colon and spleen were stimulated for 3 h at 37°C with 50 ng/ml PMA (Sigma-Aldrich) and 1 μM ionomycin (Sigma-Aldrich) in the presence of protein transport inhibitor cocktail (eBioscience) in complete RPMI 1640 supplemented with 10% FBS (Thermo Fisher Scientific). Extracellular surface antigen staining and intracellular transcription factor and cytokine staining were performed as described above.

### IL2 Experiments

#### IL2 injections

For scATAC-seq of IL2 treated Tregs, 6-week-old *Foxp3*^IRES-GFP^ mice were injected intravenously with 10 μg of recombinant mouse IL2 (PeproTech, 212-12) or PBS vehicle control (in 110 μl volume). Mice were euthanized exactly 2 hours after injection for isolation and processing of splenic Tregs, as described in previous sections.

#### pSTAT5 Flow Cytometry

For *ex vivo* quantification of STAT5 phosphorylation, after mechanical dissociation and ACK red blood cell lysis, single-cell suspensions of spleens from 10-week-old *Foxp3*^Thy^^1.1^ mice were stimulated for 15 minutes at 37°C with the indicated concentration of recombinant mouse IL2 (PeproTech, 212-12) in serum-free RPMI 1640. Stimulation was quenched by cell fixation at a final concentration of 2% PFA on ice for 30 minutes. Afterwards, cells were washed twice with ice-cold PBS and permeabilized with pre-chilled 90% methanol on ice for 30 minutes. Cells were washed and stained with pSTAT5 (Tyr694) antibody (1:20, BioLegend) at room temperature for 40 minutes, at which point, surface marker antibodies were added to samples for additional 25 minutes staining, prior to washing, resuspension, and flow cytometry data collection. Cells were distinguished as aTreg vs rTreg using CD44 and CD62L staining.

### CRISPR/Cas9 RNP Editing and Transfer

We adapted a previous protocol for gene editing using CRISPR/Cas9 ribonucleoprotein (RNP) complex nucleofection ^103^ for application to primary mouse Tregs. We isolated CD4 T cells from pooled spleen and lymph nodes of 6-8-week-old *Foxp3*^IRES-GFP^ mice using magnetic negative selection using the Dynabeads Untouched Mouse CD4 Cells Kit (ThermoFisher Scientific, cat#11415D), resting cells for at least 30 min at 4°C prior to electroporation. We freshly prepared gRNA duplexes by combining crRNA (pre-designed validated crRNA, IDT) and tracrRNA oligos (IDT) at equimolar concentrations to a final duplex concentration of 40 uM, heating the duplex at 95°C for 5 min in a PCR thermocycler and cooling down at RT for 15 minutes before keeping on ice. For every reaction, we used 12x10^6^ CD4 T cells, washed with Ca/Mg free PBS and spun down at 200G for 10 min at RT to remove FBS. During this spin, immediately prior to nucleofection, we assembled RNP complexes by thoroughly mixing 2.25 ul of 5 ug/ul TrueCut Cas9 Protein V2 (Thermo Fisher) and 5ul 40uM gRNA duplex and incubating at RT for 15 minutes. For each target, we used a pool of two separate RNPs (assembled separately) carrying different gRNAs duplexes. Cells were resuspended in 100 ul of Electroporation Solution P4 (Lonza) with the pool of both targeting RNPs or with RNPs carrying control non-targeting gRNAs for 2 min at RT and transferred into Large Lonza cuvettes for nucleofection with a Lonza Amaxa 4D Nucleofector (Program DS137). 150 ul of pre-warmed complete T cell medium (RPMI-1640, 10% FBS, supplemented with L-Glutamine, NEAA, beta-mercaptoethanol, and 10mM HEPES) with pre-equilibrated cytokine (10ng/mL recombinant mouse IL7 and 2000 IU/m recombinant human IL2) was added immediately afterwards and cells were rested for 1h at 37°C. Cells from each cuvette reaction were subsequently transferred to separate wells of a 12 well plate containing 1mL of pre-warmed complete medium, adding another 750 ul of media used to washing residual cells from cuvettes. After 2 additional hours of rest, cells were expanded with anti-CD3/anti-CD28 (Thermo Fisher) beads at a 1:1 ratio for 3 days. All cells in culture were transferred into separate Treg-depleted *Foxp3*^DTR^ hosts, injected with DT (Sigma) at 20 ng/g mouse body weight for two consecutive days prior to transfer and again on the day following transfer. Cells were parked *in vivo* for 1 week prior to sorting 10,000 GFP+ Tregs from pooled spleen and lymph nodes for bulk ATAC-seq. 3 biological replicates were used for both targeting and non-targeting conditions.

### Bulk ATAC-seq library preparation

Bulk ATAC-seq libraries were prepared using the ImmGen ATAC-seq protocol as described in ^29^.

### scATAC-seq library preparation

For non-hashtagged experiments, nuclei isolation, transposition, GEM generation, and library construction targeting capture of 10000 cells were carried out as detailed in the Chromium Next GEM Single Cell ATAC manual (10x Genomics). Libraries were pooled and sequenced on an Illumina NovaSeq 6000 to a final median depth of approximately 20-30,000 paired-end reads per cell. Sequencing data were converted to fastq files, aligned to the mm10 reference genome, and quantified per cell using Cell Ranger ATAC software (10x Genomics, v1.2).

### ASAP-seq

For experiments with multiple conditions per scATAC run, we hashtagged cells using a modification of the ASAP-seq strategy^78^ for low cell input primary cell samples. Before sorting, cells were hashtagged with mouse TotalSeqA DNA-barcoded hashtag antibodies at the same time as staining with fluorophore-conjugated antibodies (BioLegend). Hashtags used in each experiment are provided in Table S1. Cells were sorted into DMEM + 5% FCS in DNA Lo-Bind tubes (Eppendorf, cat # 022431021). After spinning down for 5 min at 500g in a refrigerated centrifuge at 4°C, cells were resuspended in 100 μl chilled 0.1x Omni Lysis buffer (1x Omni Lysis buffer (10mM Tris-HCl, 10mM NaCl, 3 mM MgCl_2,_ 0.1% Tween-20, 0.1% NP40 substitute/IGEPAL, 0.01% Digitonin, 1% BSA in nuclease free water) diluted 1:10 in Wash/Lysis Dilution Buffer (10mM Tris-HCl, 10mM NaCl, 3 mM MgCl_2,_ 1% BSA in nuclease free water)), gently mixed by pipetting and incubated on ice for 6.5 min. Following lysis, 100 μl chilled wash buffer was added and gently mixed by pipetting. Cells were spun down for 5 min at 500g at 4°C, all but 5 μl of supernatant was removed, and 45 μl of chilled 1x nuclei buffer (10x Genomics) was added without mixing. After one more centrifugation step at 500g, 4°C for 5 min, supernatant was removed, and samples were resuspended in 7ul 1x nuclei buffer for cell counting and input into transposition, barcoding, and library preparation according to the Chromium Next GEM Single Cell ATAC manual (10x Genomics).

Modifications to the original 10X protocol were made as described in the original ASAP-seq publication and as detailed at https://citeseq.files.wordpress.com/2020/09/asap_protocol_20200908.pdf. Briefly, 0.5 μl of 1uM BOA bridge oligo was spiked into the barcoding reaction. During GEM incubation, an additional 5 min incubation at 40°C was added to the beginning of the protocol. 43.5 instead of 40.5 μl of Elution Solution I was added during silane bead elution to recover 43 μl. 40 μl was used for SPRI clean up as indicated in the protocol, while 3 μl was set aside. During SPRI cleanup, the supernatant was saved. The bead bound fraction was processed as in the protocol, while for the supernatant fraction, 32 μl SPRI was added for 5 min. Beads were collected on a magnet, washed twice with 80% ethanol, and eluted in 42 μl EB. This 42 μl was combined with the 3 μl set aside from the previous step as input into the HTO indexing reaction. HTO Indexing PCR was run with partial P5 and indexed Rpxx primers (https://citeseq.files.wordpress.com/2020/09/asap_protocol_20200908.pdf) as: 95°C 3 min, 12-14 cycles of (95°C 20 sec, 60°C 30 sec, 72°C 20 sec), 72°C 5 min. The PCR product was cleaned up with 1.6X SPRI purification for quantification and sequencing alongside ATAC libraries.

### scRNA-seq Library Preparation

We sorted GFP+ (WT or KO) or Thy1.1+ (WT) Tregs and reporter negative Tconv from spleens and colonic lamina propria of heterozygote female mice for scRNA-seq. Cell encapsulation and library generation for scRNA-seq was carried out using the Chromium Single Cell 5′ v2 and V(D)J platform with Feature Barcoding (10x Genomics). Data were processed using the standard CellRanger pipeline (10x Genomics). Cells from each condition were hashed with DNA-coded TotalSeqC Hashtag and ADT antibodies (BioLegend). Complete description of hashtag information is provided in Table S1.

### scATAC-seq preprocessing and visualization

Data analysis was performed using Signac v1.4^124^. For quality control, only cells with at least 1-4x10^3^ fragments per cell (depending on sequencing depth of experiment), greater than 50 percent reads in peaks (relaxed to 30 percent in colon samples due to presence of colon-specific peaks), TSS enrichment score greater than 2, nucleosome signal less than 10, and ratio of blacklist-region reads less than 0.05 were retained for further analysis. See Table S1 for quality-control metrics for all datasets used in this study. Putative doublets identified by ArchR v1.0.1^49^ and non-Treg, non-Tconv contaminant cells were also removed. We used the latent semantic indexing approach as previously described^125, 126^. Binarized count matrices were normalized using the term frequency-inverse document frequency (TF-IDF) transformation and reduced in dimensionality by singular value decomposition (SVD). As the first component was highly correlated with sequencing depth, SVD components 2-30 were used to generate a shared nearest neighbor (SNN) graph for clustering and as input into UMAP^127^ with cosine distance metric for visualization. For visualization of B6/Cast F1 datasets, we created 2 features per OCR, one corresponding to B6-assigned reads and one corresponding to Cast-specific reads, using both versions as independent features for visualization and clustering.

### scATAC-seq Analysis

#### Hashtag counts + assignments

Hashtag processing followed the original recommendations of the ASAP-seq paper^78^, using asap_to_kite (https://github.com/caleblareau/asap_to_kite) to process FASTQs files for downstream quantification by the bustools and kite workflows^128, 129^. We used the HTODemux()^130^ function in the Seurat package (v4.0.2)^131^ to remove doublets and call hashtag identities.

#### Peak Sets

To enable comparisons across conditions and datasets, we used a common set of open chromatin regions throughout the study. After initially processing data mapped to a set of pan-immune cell OCRs defined by the Immgen consortium^29^, we used iterative sub-clustering at increasing resolution, guided by the silhouette score, to group cells into “metacells” with median size of 350 cells each. We used Archr v1.0.1^49^ and MACS2 v2.2.7.1^132^ to call fixed-width peaks of 250bp within each of these subgroups of cells, merged into a final set of high-resolution peaks. We compared *de novo* peak calls with peaks from the Immgen *cis*-regulatory atlas, keeping only those Immgen peaks that overlapped at least one peak in the *de novo* merged peak set. In cases where new Treg-specific peaks overlapped with an existing Immgen peak, we retained the Immgen version for cross-reference compatibility. In cases where a Treg peak did not match an Immgen peak, it was added to the reference peak set. In total, our final Immgen+Treg reference OCR set consisted of 216419 peaks and is provided in Table S2. Unless otherwise stated, this OCR set was used for all datasets throughout the study, with two exceptions. In the spleen and colon Treg comparisons, we re-called peaks in colon Tregs and added any missing regions to the reference peak set for that analysis. In the B6/Cast F1 analysis, to avoid missing Cast-specific peaks, we called peaks on Cast allele-specific reads and added any new regions to the peak set for F1 analyses. New peaks were also called to capture any additional IL2 response elements. All additional peaks are provided in Table S10.

#### Motif Accessibility Analysis

Bias-corrected relative motif accessibility was calculated using chromVAR^133^. We used motifmatchr (https://github.com/GreenleafLab/motifmatchr) to scan OCRs in our refence set from the curated set of mouse motif PWMs from the Buenrostro lab (https://github.com/buenrostrolab/chromVARmotifs/tree/master/data/mouse_pwms_v2.rda) or for all individual human and mouse motifs corresponding to ENCODE motif archetypes (v1.0)^51^ (https://github.com/jvierstra/motif-clustering), creating a merged MEME file from all included motif databases. Individual motif models and their assignment to archetypes were obtained from the information provided in ^51^. For display of motif accessibility at the level of archetypes, accessibility of individual motif models per archetype were averaged.

#### Number archetype motifs per OCR

To calculate the number of unique motif archetypes per OCR, we defined an OCR to be positive for a motif archetype if at least 50% of the motifs belonging to the archetype of interest had a match in that OCR.

#### Motif Variability

Motif Variability was calculated as the standard deviation of the chromVAR bias-corrected motif accessibility scores.

#### Gene Scores

Gene scores were calculated with Archr v1.0.1, using an exponentially weighted function that accounts for the activity of distal OCRs in a distance-dependent manner^49^ and provides an approximate proxy for gene expression. Gene modules scores were calculated using the AddModuleScore() function on gene scores in Seurat, using signatures corresponding to RORγ or Helios Tregs defined from ^134^.

#### OCR Signature Generation

OCR signatures distinguishing aTreg and rTreg populations for annotation in Figure 1 were derived from tables provided in ^47^, using OCRs with a greater than 2-fold increase in accessibility in aTregs. OCR signatures (i.e. aTreg vs rTreg, Treg vs Tconv) derived from the scATAC-seq data used OCRs with |average log_2F_C| > 0.25 using the FoldChange() function in Signac.

#### OCR Signature Relative Accessibility

Relative accessibility of OCR sets, including topics, signature OCRs, etc, was calculated using the chromVAR computeDeviations() function^133^.

#### Pseudobulk Track Visualization

To visualize pseudobulk profiles, BAM files containing reads for each group of cells were extracted using Sinto (https://github.com/timoast/sinto), shifted to account for Tn5 cut-sites, and converted to bigwigs using deeptools^135^ for display in the Integrative Genomics Viewer^136^ or the WashU Epigenome Browser^137^.

#### OCR Variance

OCR variance was calculated from log-transformed, quantile-normalized metacell counts from the scATAC-seq data as described in the peak calling section. Variance was calculated across these normalized metacell values using the rowVars() function in the matrixStats package (https://github.com/HenrikBengtsson/matrixStats).

#### IL2 Cell Matching

To pair cells with the most similar chromatin cell states in the IL2 treated vs untreated conditions, we used Harmony ^89^ to remove the IL2 effect from the LSI embedding of the scATAC-seq data. In the treatment effect-removed LSI embedding, closest cells were identified as the nearest neighbor on the k-nearest-neighbor graph constructed from Harmony corrected LSI dimensions 2 to 30 using the FindNeighbors() function in Seurat. A subsample of nearest neighbor pairs was visualized on the original UMAP embedding, coloring the visualization by the starting cell state (rTreg vs aTreg) of the untreated cell in each pair.

### OCR UMAP Visualization

To visualize the global structure of OCR usage across Treg and Tconv single cells, we used latent semantic indexing of OCRs instead of cells. After running TF-IDF normalization of the Cell x OCR matrix, we reduced dimensionality with SVD, retaining components 2-30. We used cosine similarity between individual OCRs as input into UMAP^127^ to generate the final visualization. SVD components 2-30 were used for Leiden clustering^138^ of OCRs to identify the TSS-enriched cluster. To visualize subpopulation-specific OCR signatures, per OCR |average log_2F_C| calculated using the FoldChange() function in Signac was overlaid onto the OCR UMAP.

### TF Binding and Chromatin Accessibility Data (from repository)

FASTQ files for published TF-binding and chromatin accessibility datasets were downloaded from the NIH Sequence Read Archive. Reads were trimmed using Trimgalore v0.6.6 (https://github.com/FelixKrueger/TrimGalore), aligned using Bowtie2^139^, filtered to retain high quality, singly-mapped reads with Samtools^140^, and duplicates removed using Picard (http://broadinstitute.github.io/picard/). Peaks were called using MACS2^132^. For TF-binding datasets, peaks were called in comparison to corresponding IgG and/or TF knockout controls. Peaks with irreproducible discovery rate^141^ < 0.05 were kept as high-quality binding annotations. In cases where binding sites were provided in the reference publication, we used the provided sites, lifting over coordinates to the mm10 genome when necessary. For FoxP3 binding analysis, we used a previously defined set of robust FoxP3 binding sites^29, 39^, derived from intersecting FoxP3-bound regions from 2 different datasets^28, 58^.

### OCR-Gene Correlation Analysis

We used FigR v0.1.0 to determine OCR to gene correlations^71^. We used our splenic Treg scATAC-seq and splenic WT Treg profiles from the FoxP3 KO/WT heterozygote female scRNA-seq dataset for paired analysis. Briefly, scRNA-seq and scATAC-seq profiles were first integrated using CCA^126^ on variable genes (in RNA) and gene scores (in ATAC). Integrated embeddings were used to match cells across datasets using the scOptMatch algorithm. After pairing, correlations between gene expression and OCR accessibility for regions within 100kb of each TSS were determined and compared to a background null model. For gene-OCR correlations, we kept OCRs with correlation p < 0.05 and additionally annotated any other OCRs within 15kb of the gene TSS that did not meet these significance criteria.

### B6/Cast F1 Allele-Specific Read Alignment

We adapted the diploid pseudogenome alignment strategy implemented in the lapels/suspenders pipeline^97^ and used in recent studies^47, 95, 96, 142^ for allele-specific mapping. We obtained B6 and Cast pseudogenomes, MOD files, and variant vcf files from the UNC collaborative cross project (http://csbio.unc.edu/CCstatus/index.py?run=Pseudo) and Mouse Genome Project^91^. Reads were aligned to both B6 and Cast pseudogenomes, shifted to a common set of B6-based reference coordinates, and assigned to an allele of origin based on which allele has stronger mapping. Non-specific reads with equally strong mapping to both alleles were randomly split in half into B6 and Cast groups to obtain the final reads used for allele-specific analyses. Allele-specific BAMs were converted to fragments files using Sinto for input into downstream scATAC analysis.

### Ensemble Topic Modeling

Further details on ensemble topic modeling are provided in Computational Note 1.

#### TSS vs Distal OCRs

We defined TSS OCRs as those OCRs which overlapped an annotated TSS as retrieved from UCSC mm10 annotations (http://hgdownload.cse.ucsc.edu/goldenPath/mm10/database/refFlat.txt.gz). Distal OCRs were defined as those OCRs that did not overlap annotated TSS. Because OCRs not overlapping TSS in the TSS-predominant cluster (Fig S2A) on the OCR UMAP had similar patterns of accessibility to TSS-overlapping OCRs and mapped closer to TSS than OCRs outside of this cluster, this cluster likely contained promoter OCRs not within 250bp of the TSS. Thus, any OCRs in the TSS cluster were removed from the distal OCR set for analysis.

#### Topic Motif Enrichment

To calculate enrichment of motifs within topic OCRs, we used a permutation testing framework. We compared the number of observed motif matches within each topic (separately by TSS or distal designations) to the number of matches among a set of 100 background OCRs matched for GC content and accessibility (chosen using the chromVAR getBackgroundPeaks() function). Significance was assessed using a two-sided Z test, with Benjamini-Hochberg false discovery rate correction. We kept motif enrichments with FDR < 1x10^-10^. To filter out noise, we considered only motifs of TFs expressed in Tregs and present in at least 3% percent of queried OCRs (TSS or distal) in that topic.

#### Topic OCR enrichment (Tissue)

We defined a set of pan-tissue Treg OCRs as OCRs with at least a 2-fold increase in accessibility relative to spleen Tregs across all comparisons of Tregs from muscle, colon, and visceral adipose tissue in a previously published dataset^16^. We then used the same permutation framework described above to look for enrichment of these OCRs within each topic.

#### TF Motif, Binding Overlap Analysis

To quantify significance of overlaps in enrichments of TF binding sites and motif within topics, we first computed separate enrichments for TF and matched motifs in each topic using the permutation framework described above, keeping enrichments with FDR < 1x10^-10^ in both comparisons. Overlap between the two sets of enrichments was quantified using a binomial test where success was defined as concordance (for presence or absence of enrichment).

#### Topic Variance Explained

We used the reconstruction error from a linear regression to calculate the variance explained by each topic. For each topic, we fit a linear regression model with centered, Treg metacell pseudobulk accessibility as the response variable and an indicator variable of whether or not each OCR was assigned to a topic as the predictor variable (with 0 intercept). The proportion of total variance in the metacell accessibility explained by the fit regression model was used to estimate the percentage of variance in accessibility explained by each topic. Note that this will be an underestimate in actual variance explained, given that 1) this uses a linear regression framework, 2) looks at variance across single cells collapsed to the metacell level, and 3) uses binarized topic OCR assignments. However, the approach is still a useful approximation for comparing relative variance explained across different conditions.

#### Differential Topic Accessibility

To calculate differential topic accessibility, we calculated the log_2 F_old Change in average accessibility of OCRs from each topic between the two conditions of interest. To assess significance, we used a two-sided t-test between the log transformed values from each condition, correcting p values for multiple hypothesis testing using the Benjamini-Hochberg false discovery rate procedure.

#### GREAT Analysis

To link regulatory regions in each topic to gene sets, we used GREAT ^76^, as implemented by the rGREAT package ^143^. We used the rGREAT::great() function with default parameters and ‘txdb:mm10’ as TSS source to look for enrichment of gene ontology term annotations in topic regions. For interpretability and to avoid an unreasonably large number of terms, we restricted enrichment to gene ontology terms from the Biological Process category, with “immune system process”, “response to stimulus”, or “signaling” annotations. Enrichments with p-value < 1x10^-10^_, f_old enrichment > 4, fraction genes in gene set > 50%, and observed region hits > 20 were kept for display.

#### IL2 Cell State Assignment

To assign cells to rTreg, aTreg, or rorTreg state in the IL2 scATAC-seq analysis, we first computed per-cell relative accessibility (chromVAR scores) of aTreg vs rTreg and aTreg vs rorTreg OCR signatures. We then fit a 3-component gaussian mixture model to the matrix of per-cell signature accessibility scores to classify cells as rTreg, aTreg, or rorTreg.

##### HOMER motif enrichment

We ran motif enrichment in OCRs from each cluster of differential IL2 responses using HOMER ^144^. We used the findMotifsGenome.pl with a background set of all Treg OCRs defined in this study, motif length of 10, scan size of ‘given’, and -S set to 25. Top motifs with p value < 10^^-5^ and at least 5% hits in each OCR cluster were kept as significant.

### Topic AME

Further details on the approach for quantifying topic-specific AMEs are provided in Computational Note 2.

### TF KO Analysis

In the case of *Foxp3-cre*×*Gata3^fl/fl^* and *Foxp3-cre*×*Gata3^+/+^*littermate comparisons, cells corresponding to each genotype were aggregated into pseudobulks, filtering out OCRs with fewer than a mean of 5 reads across samples. In the case of c-Maf knockout data, we used bulk ATAC-seq counts from ^98^ intersected with our reference peak set, once again filtering out OCRs with fewer a mean of 5 reads across samples. Samples were quantile normalized prior to computing the fold change between KO and WT conditions. To narrow in on likely direct effects, we identified distal (non-TSS) OCRs with a greater than two-fold decrease in accessibility in the knockout vs wildtype that contained GATA or MAF archetype motifs, respectively. We then computed enrichment of each of these OCR sets within topic OCRs using a permutation test versus matched background OCRs as described above. For Smarcc1 analysis, we used DESeq2 ^145^ to calculate differential accessibility between bulk ATAC-seq samples from *Smarcc1* edited vs control cells (3 biological replicates for each condition). Distal (non-TSS) OCRs with mean base accessibility greater than 5 and loss of accessibility in the *Smarcc1* edited condition with p value < 0.05 were selected as Smarcc1-dependent OCRs. We computed topic OCR enrichment using a permutation test within this OCR set as described for other TF KO analyses.

### FoxP3 heterozygous female analysis

#### Differential OCRs

We calculated differential accessibility using a logistic regression per OCR with number of fragments per cell included as a latent variable. OCRs with average |log_2 F_old Change| > 0.1 and p value < 0.05 were designated as differential. To avoid effects driven by cell composition, we computed differentials separately subclustered on cells corresponding to rTregs and aTregs from each genotype, based on chromVAR scores of aTreg vs rTreg signature OCRs. For FoxP3-independent OCR comparisons, we compared rTreg KO cells with Tconv.

#### Differential accessibility of motifs within each topic

To calculate differential accessibility of motifs within each topic, we calculated the log_2 F_old Change in average accessibility of topic OCRs containing the motif of interest between FoxP3 WT and KO cells. To avoid effects driven by cell composition, we computed differentials separately between clusters corresponding to rTregs and aTregs, as per above. To assess significance, we used a two-sided t-test between the log transformed values from each condition, correcting p values for multiple hypothesis testing using the Benjamini-Hochberg false discovery rate procedure. Effects with FDR < 0.05 were displayed on the heatmap. FoxP3-independent OCR comparisons were done in the same manner but instead comparing rTreg KO cells with Tconv.

#### Local Inverse Simpson’s Index

The Local Inverse Simpson’s Index (LISI) is a measure originally developed for evaluating efficacy of batch integration methods in single cell genomics, but which provides a general metric for overlap between populations^89^. To use LISI to compare overlap between FoxP3 KO and WT vs Gata3 KO and WT cells, we first integrated scATAC profiles from both datasets into a common embedding using Seurat^126^. We computed diversity in local neighborhoods using the LISI metric from the LISI package for both FoxP3 KO and WT vs Gata3 KO and WT cells, using the LSI components 2-30 in this common, integrated embedding (https://github.com/immunogenomics/LISI). For IL2 comparisons, we used the original LSI components 2-30 for LISI computations, as these cells were already in a common embedding within the same experiment.

### scRNA-seq Analysis

scRNA-seq data analysis was performed in Seurat (v4.0.2) ^131^}. Cells with greater than 500 UMIs and fewer than 10% mitochondria reads were kept for downstream analysis, with contaminant cells removed using known marker genes. We used the HTODemux()^130^ function to remove doublets and call hashtag identities. The top 2500 variable genes and first 35 principal components were used for nearest-neighbor graph construction and UMAP visualization. Tregs were classified as RORγ+ using signatures from ^24^. Treg signature scores were calculated as the output from the AddModuleScore() function for the Treg Up signature minus the output for the Treg Down signature ^31^.

### Data Visualization

Graphs and visualizations were generated in R (v4.1.0^146^) using ggplot2 (v2.3.3.5^147^) and in Python (v3.9.7, http://www.python.org) using matplotlib v3.5.0^148^. Flow cytometry data was analyzed using FlowJo v10 (BD LifeSciences), with corresponding plots and statistical analyses done in GraphPad Prism (www.graphpad.com).

### Data Availability

Raw and processed data files have been deposited at the Gene Expression Omnibus (GEO: GSE216910 and GSM5712663).

## SUPPLEMENTARY FIGURE LEGENDS

**Figure S1:**
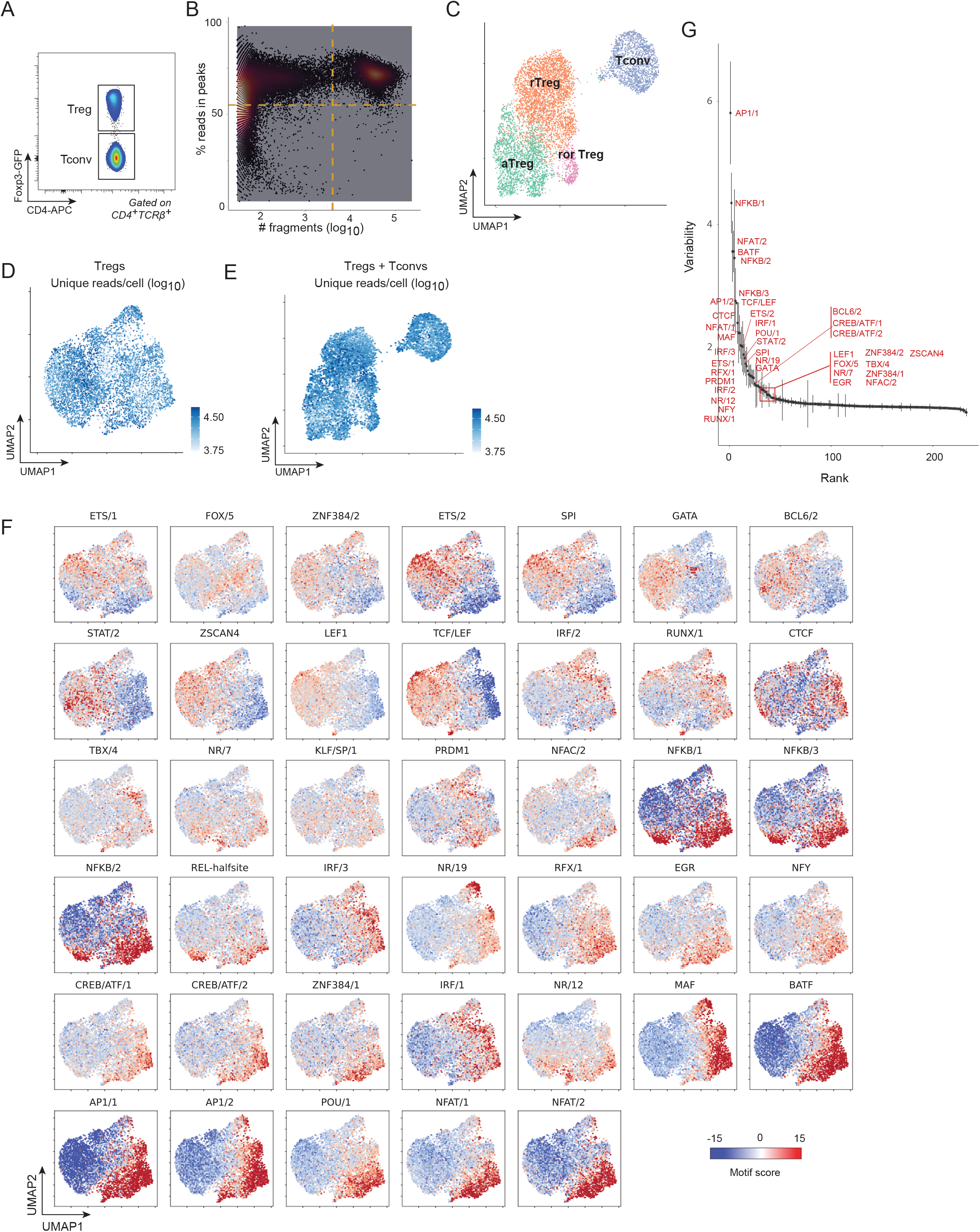
Additional analysis of Treg scATAC-seq data. A) Sort strategy for isolation of Treg and Tconv cells from *Foxp3*^IRES-GFP^ reporter mouse. B) Fraction of reads in peaks vs number of unique reads per cell plot for quality control of scATAC-seq data. Dotted lines indicate thresholds used to select cells for analysis. C) UMAP of scATAC-seq data of splenic Treg and Tconv from *Foxp3*^IRES-GFP^ reporter, colored by subpopulation annotation. D) UMAP of Treg scATAC-seq from Figure 1 colored by number of unique reads per cell. E) UMAP of Treg and Tconv scATAC-seq from (C) colored by number of unique reads per cell. F) Relative accessibility (chromVAR scores) of OCRs containing indicated TF motifs for most variable TF motifs across Treg scATAC-seq data; motifs averaged within ‘archetypes’ to reduce redundancy. Only motifs whose corresponding TF(s) are expressed in Treg cells are shown. G) Variability of motif accessibility (chromVAR scores) across Treg scATAC-seq data.

**Figure S2:**
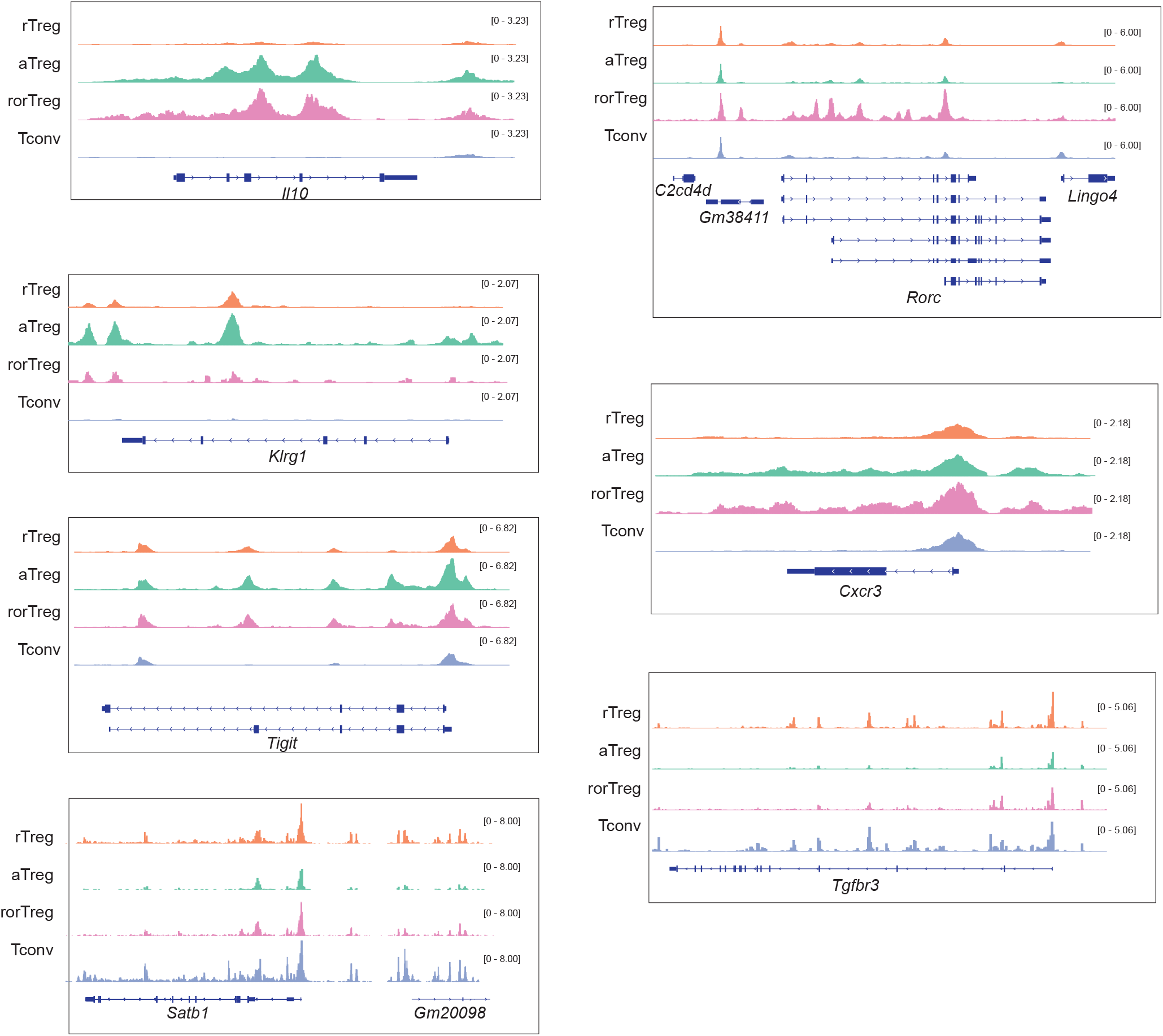
**Genome Browser Views of State-Specific Loci**. Aggregated accessibility of cells from rTreg, aTreg, rorTreg, or Tconv cell states at state-specific loci from scATAC-seq data in Figure 1.

**Figure S3:**
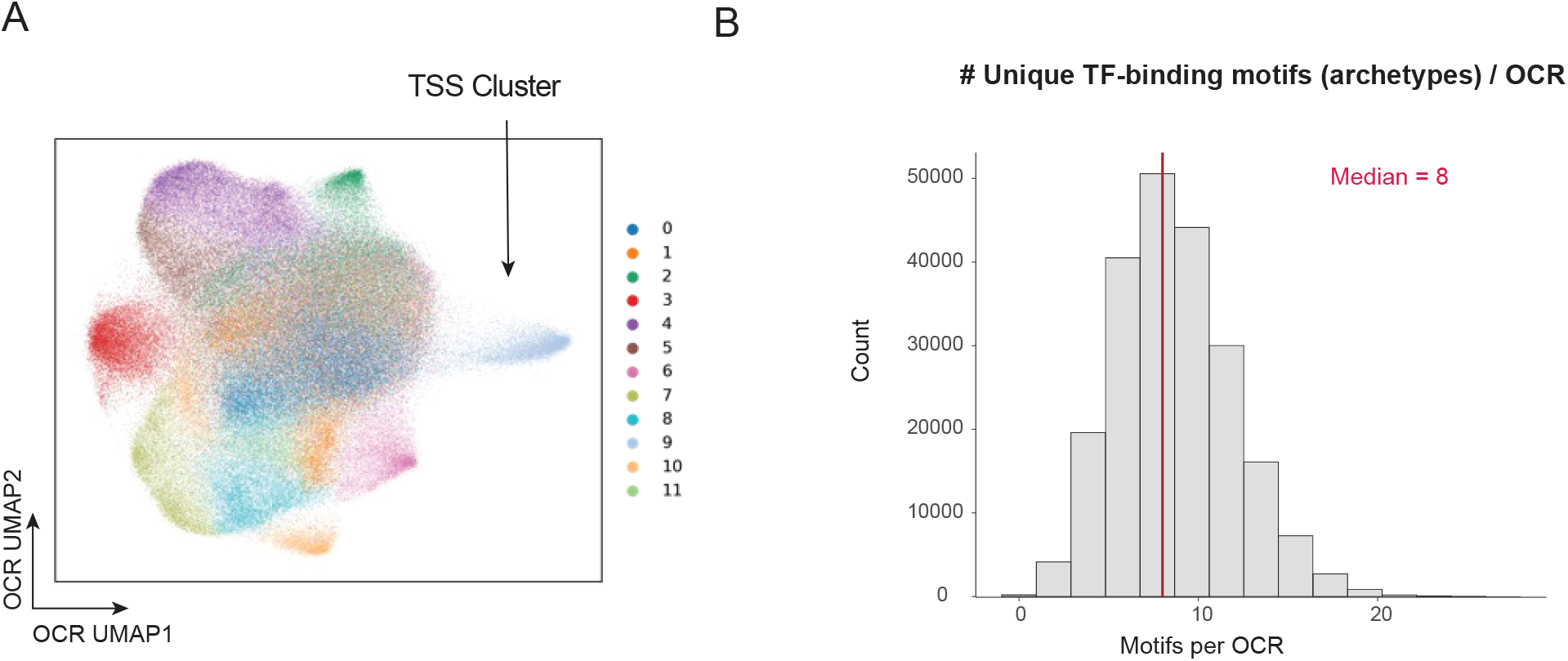
Additional analysis of OCR distributions. A) OCR UMAP from Fig 2, colored by Leiden clustering of OCRs, with cluster enriched for TSS indicated. B) Distribution of number of unique archetype motifs per OCR

**Figure S4:**
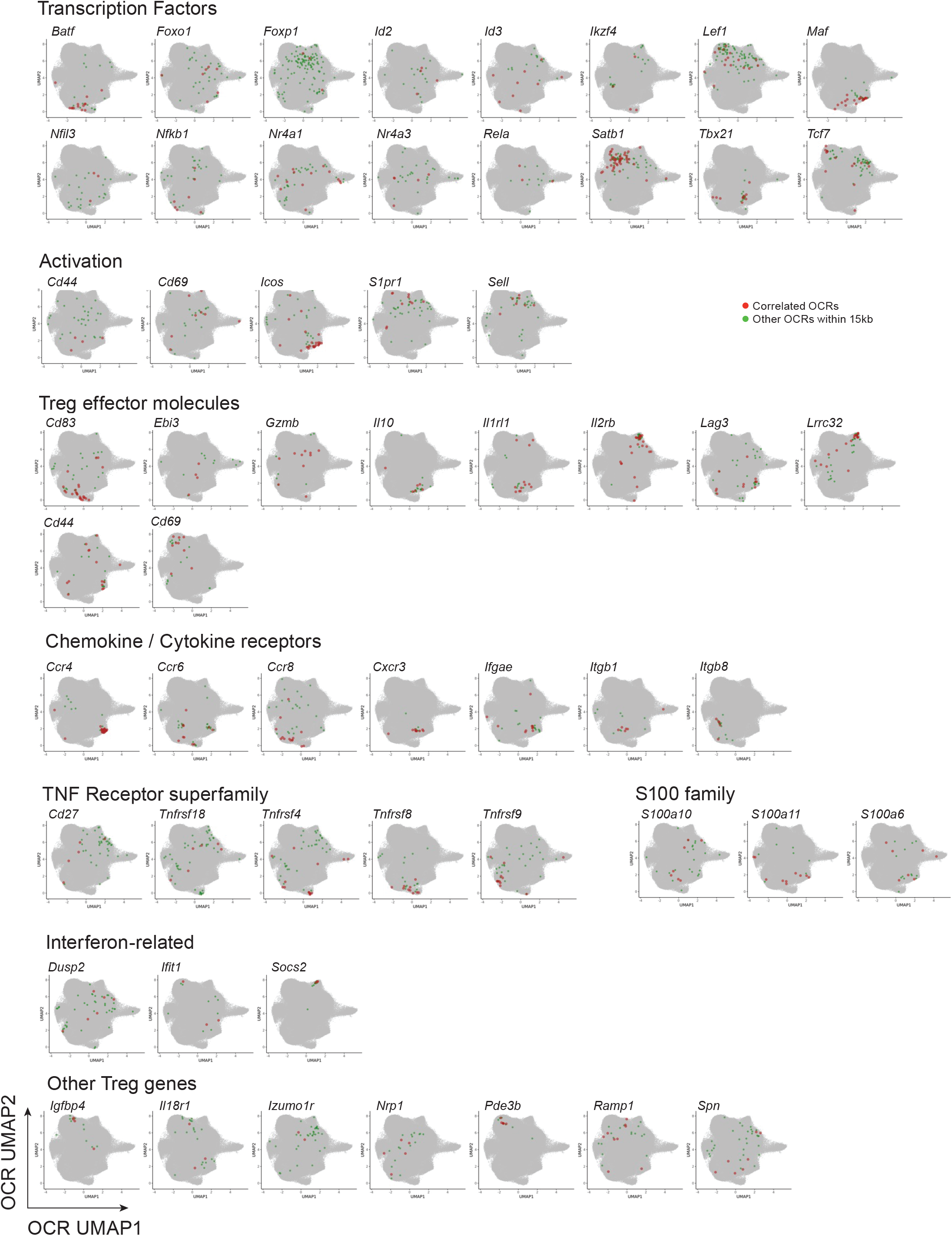
Additional Gene-OCR Correlations. Annotation on OCR UMAP from Fig 2 of OCRs with accessibility correlated (FigR p < 0.05) with expression of indicated genes (in red) or other OCRs within 15 kb of gene TSS not meeting this correlation significance threshold (in green).

**Figure S5:**
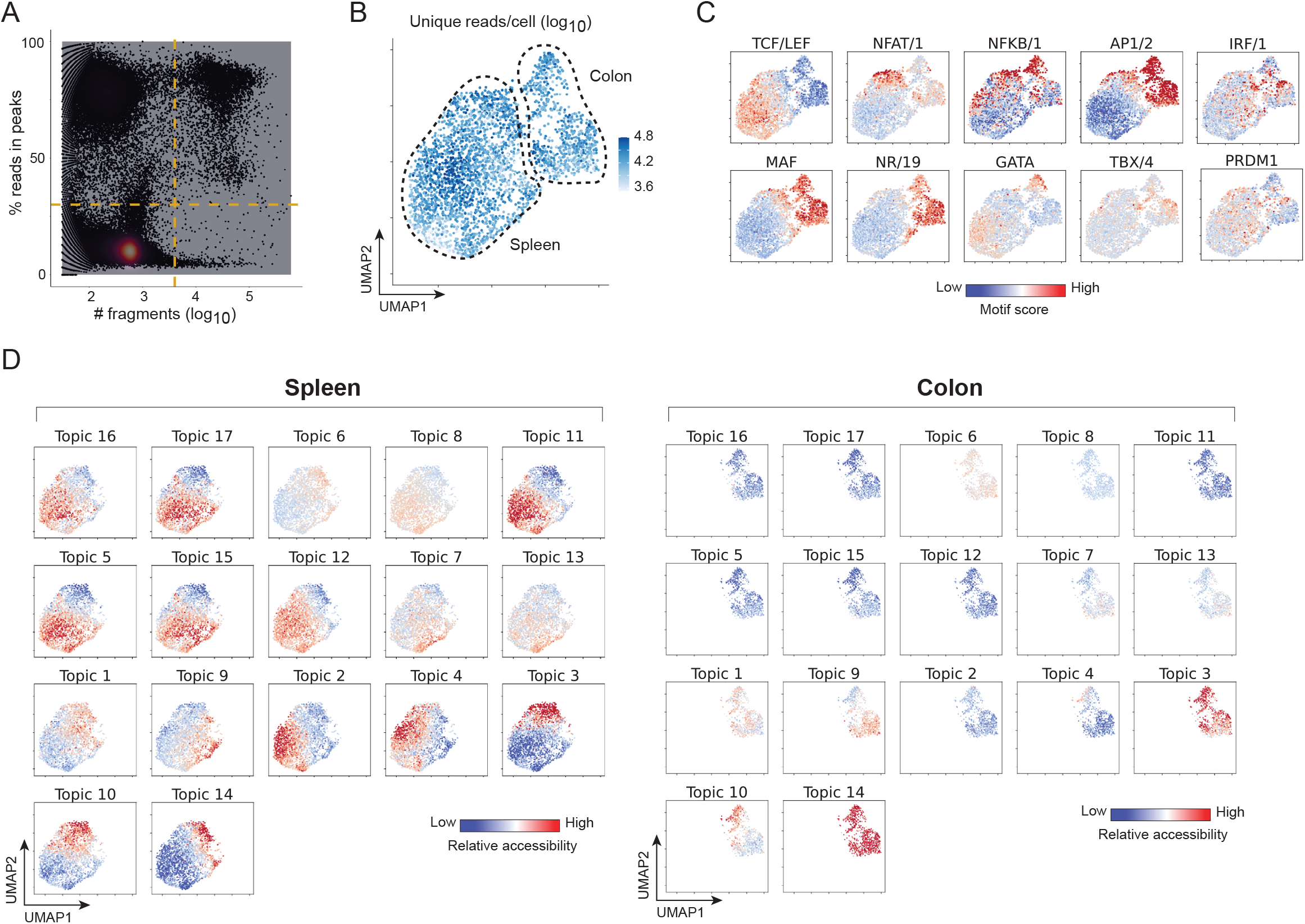
Additional comparisons of Spleen and Colon Treg scATAC profiles. A) Fraction reads in peaks vs number of unique reads per cell plot for quality control of scATAC-seq data. Dotted lines indicate thresholds used to select cells for analysis. B) UMAP of Treg scATAC-seq from Figure 4 colored by number of unique reads per cell. Dotted lines separate spleen and colon Tregs. C) Relative accessibility (chromVAR scores) of OCRs containing indicated TF motifs across spleen and colon Treg single cells visualized on scATAC UMAP from Figure 4, separated by organ. D) Relative accessibility (chromVAR scores) across spleen and colon Treg single cells of OCRs from all topics visualized on scATAC UMAP from Figure 4, separated by organ.

**Figure S6:**
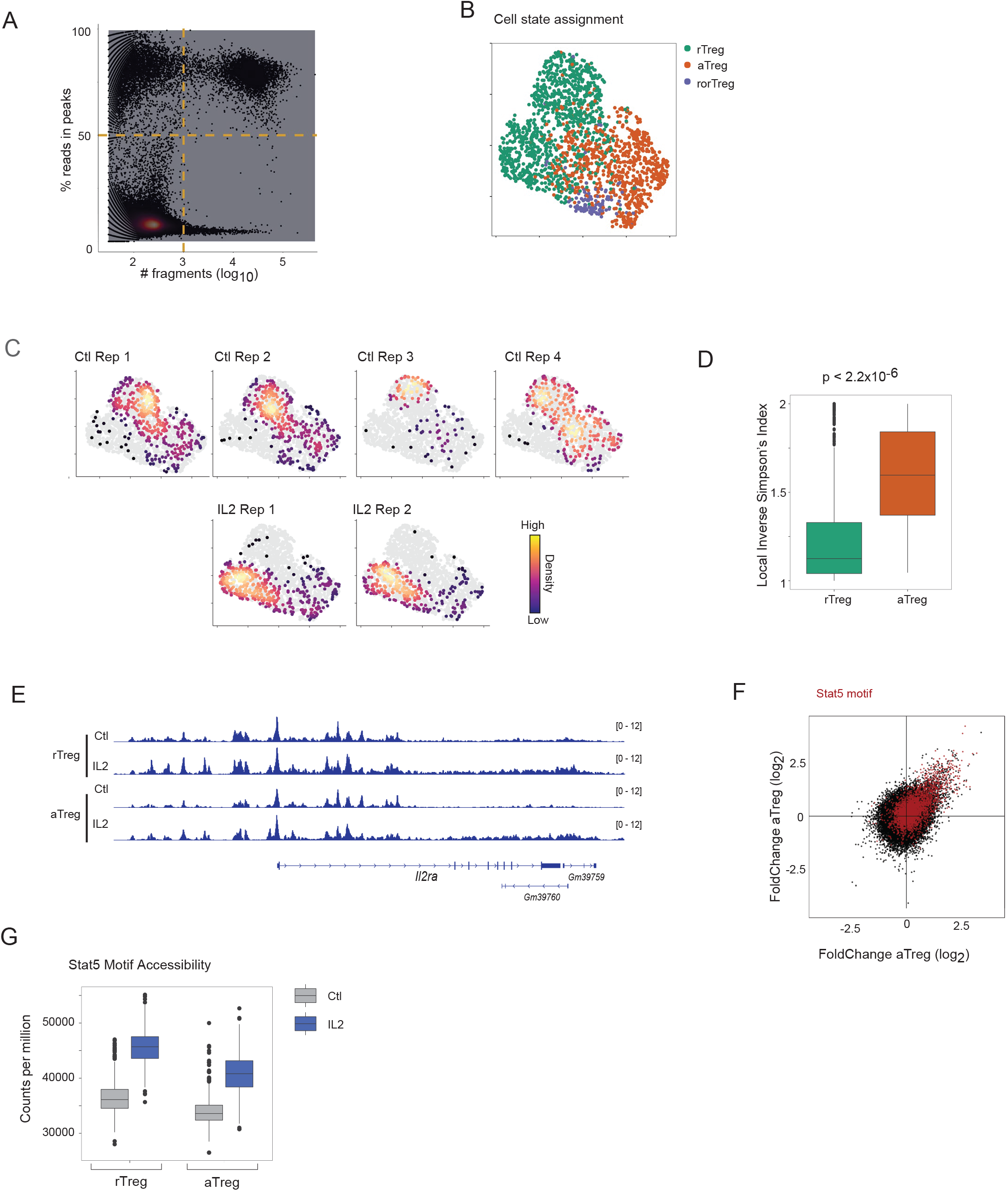
Additional analysis of IL2 scATAC-seq data. A) Fraction reads in peaks vs number of unique reads per cell plot for quality control of scATAC-seq data from Fig 5. Dotted lines indicate thresholds used to select cells for analysis. B) Classification of cells in scATAC-seq data from Fig 5 by cell state based on relative accessibility of OCR signatures distinguishing these populations derived from data in Fig 1. C) UMAP of scATAC-seq of IL2 -treated or -untreated splenic Tregs from Fig 5, colored by density of cells from each condition in each biological replicate. D) Overlap between IL2 treated and untreated cells within rTreg or aTreg pools, as measured by the Local Inverse Simpson’s Index (LISI). LISI values range from 1 for complete separation to 2 for complete overlap (p value from Wilcoxon Rank Sum Test). E) Aggregated accessibility profiles at *Il2ra* locus of IL2 treated or untreated Treg single cells from scATAC-seq data in Figure 5, separated by cell state (rTreg or aTreg). F) Fold Change-Fold Change plot showing log_2F_old Change in accessibility of each OCR after IL2 treatment in aTreg vs rTreg conditions. OCRs containing Stat5 motif are indicated in red. G) Distribution of per-cell raw accessibility (counts per million) of Stat5 motif containing OCRs in Ctl or IL2 treated cells compared across rTreg and aTreg cell states.

**Figure S7:**
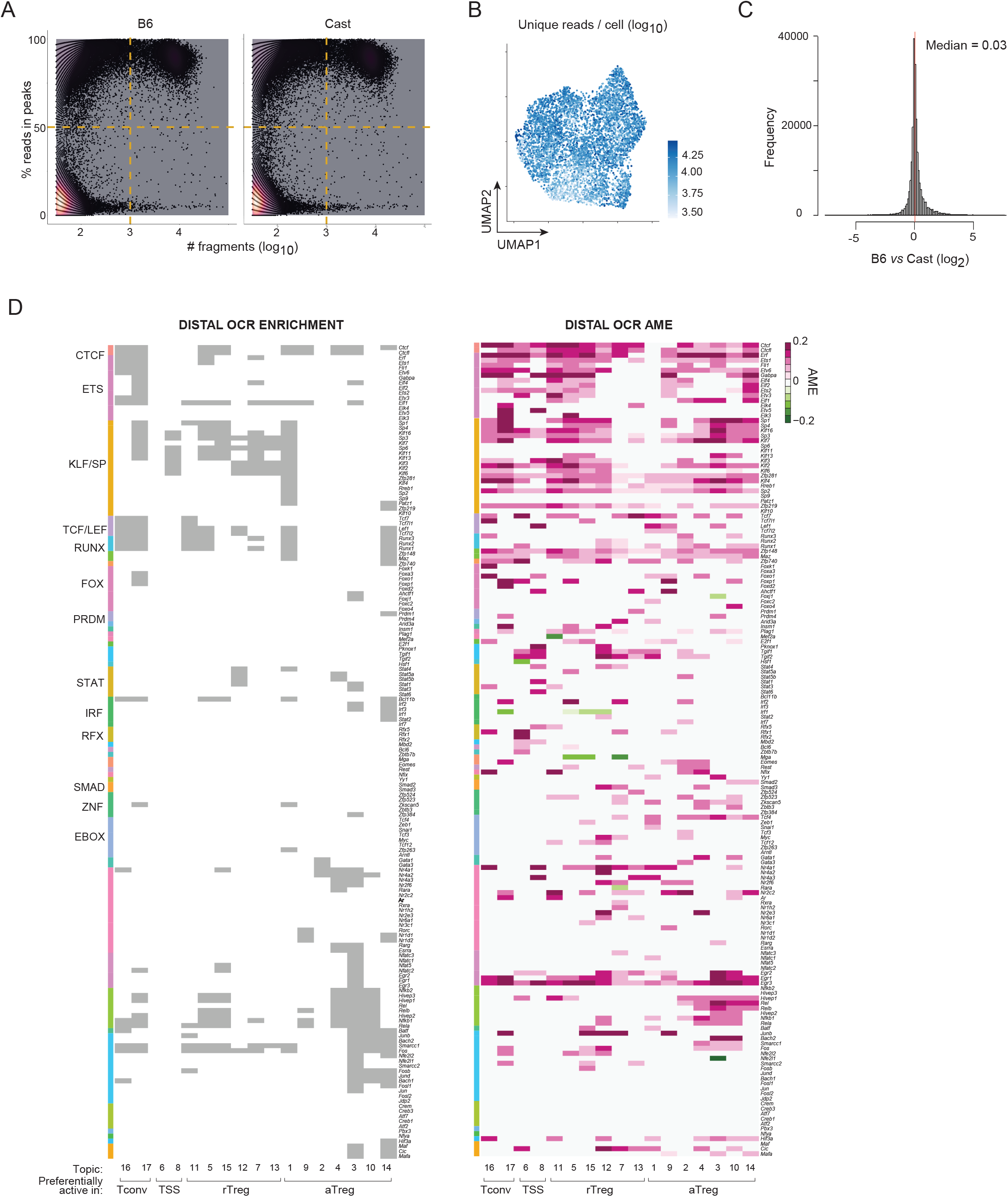
Additional F1 genetic variation analysis. A) Fraction reads in peaks vs number of unique reads per cell plot for quality control of scATAC-seq data, separated by reads assigned to B6 or Cast alleles. Dotted lines indicate thresholds used to select cells for analysis. B) UMAP of F1 Treg scATAC-seq colored by number of unique reads per cell. C) log_2 (_B6/Cast) Allelic Ratio for all OCRs aggregated across all F1 Treg single cells from Figure 6. Red line indicates median. Motif enrichment (left, grey indicates FDR<10^-10^) and significant Topic AME (right, FDR<0.10) for distal OCRs (not filtered on intersection of analyses), with heatmap order as in Figure 6B, with motifs grouped by TF family.

**Figure S8:**
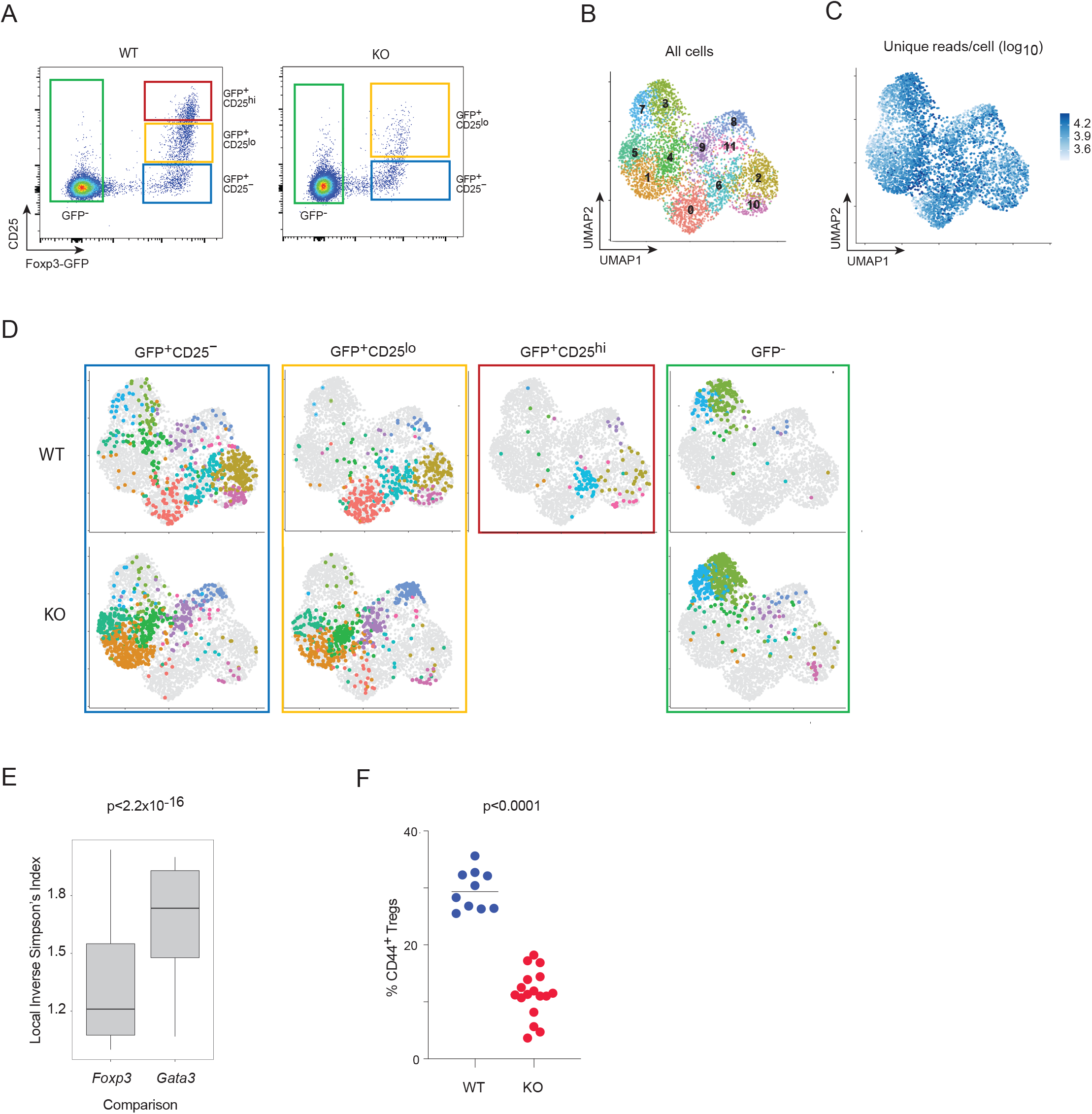
Additional analysis of FoxP3 heterozygous female scATAC-seq data. A) Sort strategy used for cell hashtagging in WT and KO FoxP3 heterozygote scATAC-seq experiment in Fig 7. B) Combined UMAP of Tconv, KO Treg, and WT Treg as in Figure 7, colored by Leiden clusters. C) scATAC-seq UMAP from Figure 7, colored by number of unique reads per cell. D) UMAP from Fig S8B separated by hashtag assignment, with colored boxes corresponding to sort gates from S8A. E) Overlap between FoxP3 KO and WT populations (left) vs between Gata3 KO and WT populations (right), as measured by the Local Inverse Simpson’s Index (LISI). LISI values range from 1 for complete separation to 2 for complete overlap (p value from Wilcoxon Rank Sum Test). F) Quantification of CD44+ Tregs from Fig 7D across biological replicates.

**Figure S9:**
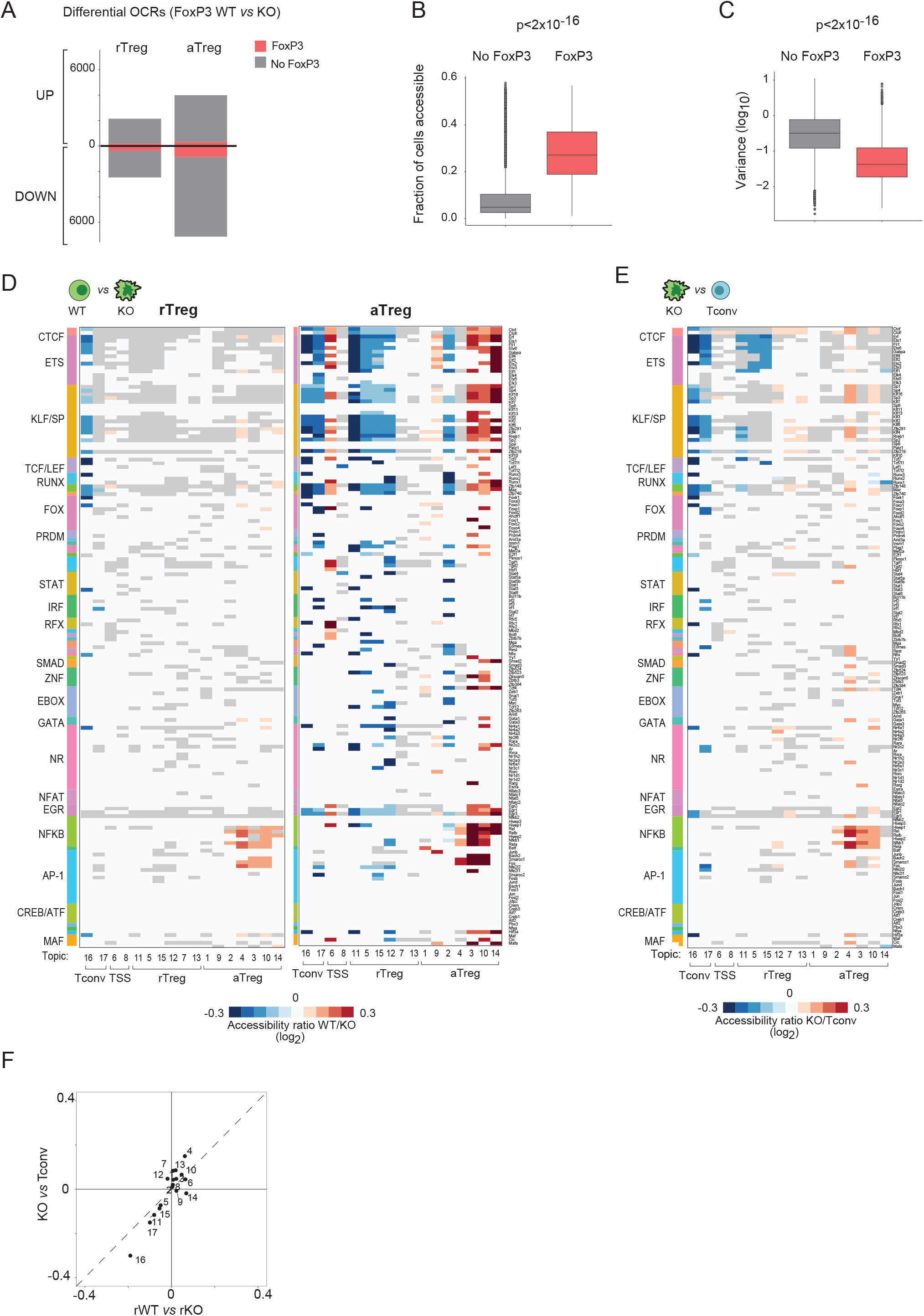
Analysis of FoxP3-bound regions and FoxP3-dependent chromatin accessibility changes. A) Number of differential (average |log2FC|>0.1, p< 0.05) OCRs in rTreg or aTreg comparisons in data from Fig 7, colored by FoxP3 binding status. B) Fraction of WT Treg cells in which OCRs are accessible based on FoxP3 binding status (p value from Wilcoxon Rank Sum Test). C) OCR variance across WT Tregs based on FoxP3 binding status (p value from Wilcoxon Rank Sum Test). D) Differential accessibility per motif in each topic (distal OCRs) between WT and KO Treg cells in rTreg or aTreg comparisons for all motif to topic connections with significant AME as in Fig S7D. Grey indicates significant topic AME in Figure S7D but no significant (FDR < 0.05) change in accessibility across FoxP3 comparisons. E) Differential accessibility per motif in each topic (distal OCRs) between KO rTreg and Tconv cells for all motif to topic connections with significant AME as in Fig S7D. Grey indicates significant topic AME in Figure S7D but no significant (FDR < 0.05) change in accessibility. F) Comparison of differential accessibility per topic in rTreg KO vs Tconv cells (y-axis) vs in rTreg WT vs KO (x-axis) cells.

**Figure S10:**
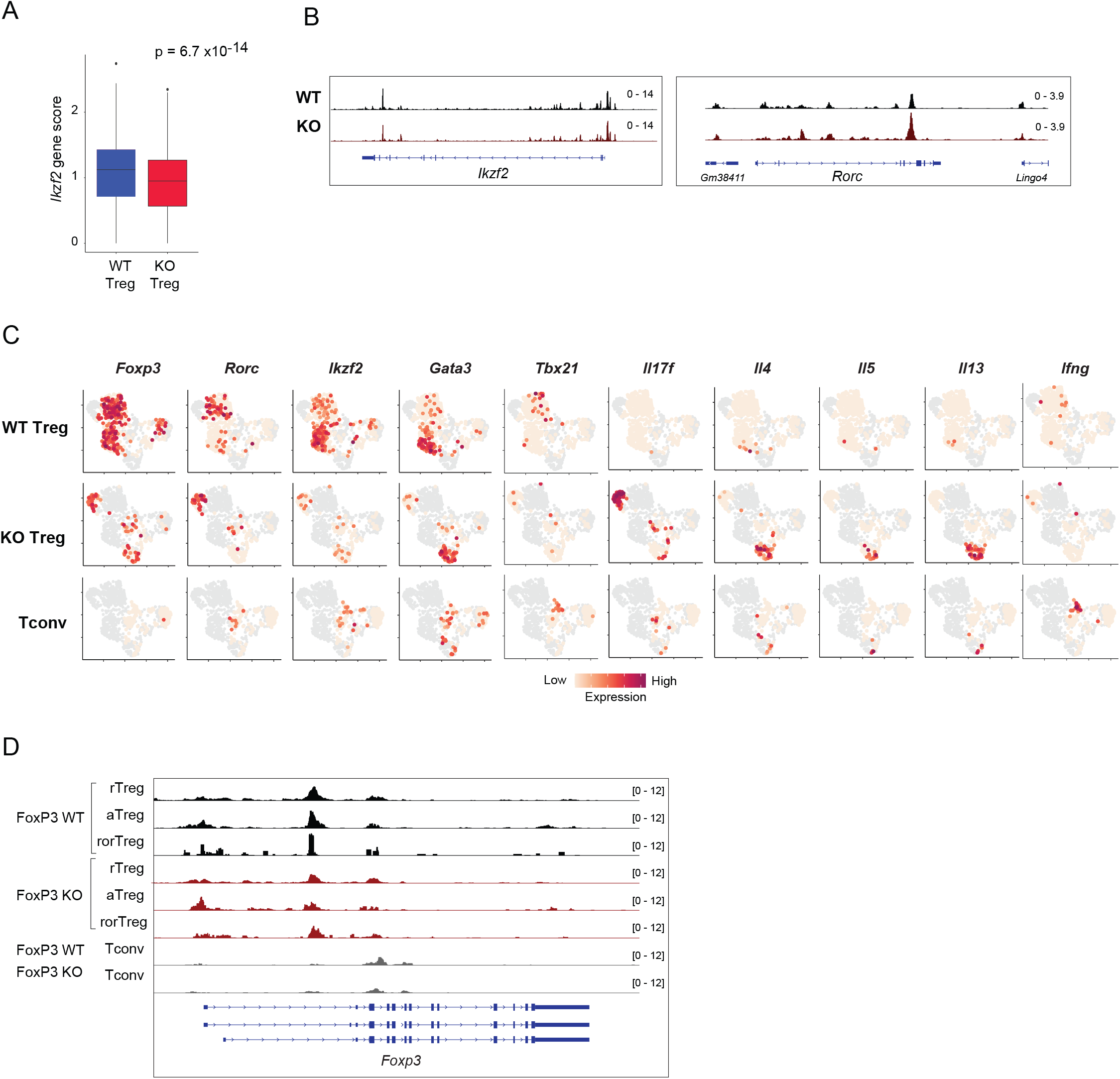
Additional analysis of FoxP3-independent RORγ+ Treg-like cells. A) *Ikzf2* gene scores in WT and KO Tregs in heterozygous female scATAC-seq data from Fig 7 (p value from Wilcoxon Rank Sum Test). B) Aggregated accessibility at *Ikzf2* and *Rorc* loci of single cells from WT or KO Tregs in heterozygous female scATAC-seq data from Fig 7. C) Expression of TF and cytokine transcripts overlaid onto colon scRNA-seq UMAP from Fig 8G. D) Aggregated accessibility at the *Foxp3* locus of single cells from WT Treg, KO Treg, and Tconv from heterozygous female scATAC-seq data from Fig 7, separated by cell state.

## SUPPLEMENTARY TABLE LEGENDS

**TABLE S1: Sample characteristic for all datasets included in this study.** A) scATAC and scRNA dataset metrics. B) Hashtags used for ASAP-seq or scRNA-seq hashtagging in each experiment.

**TABLE S2: Treg OCRs.** Information on Immgen+Treg OCRs used throughout this work, as well as additional information related to OCR UMAP and condition-specific changes in accessibility

**TABLE S3: Chromatin Accessibility and Gene Expression Signatures**. OCRs and genes used as differential signatures. In cases where external signatures were used, the provided table indicates their overlap with the reference OCRs from this study. A) aTreg vs rTreg OCRs (from ^47^). B) aTreg vs rTreg OCRs (from scATAC-seq). C) Treg vs Tconv OCRs (from scATAC-seq). D) Spleen RORγ+ Treg vs aTreg OCRs (from scATAC-seq). E) FoxP3 WT vs KO OCRs. F) Pan-tissue Treg OCRs (from ^16^). G) Gata3-(from scATAC-seq) and cMaf-dependent (from ^98^) OCRs. H) Rorγ and Helios Treg-specific gene signatures (derived from ^134^). I) Smarcc1-dependent OCRs.

**TABLE S4: TF-Binding Table.** Overlap of TF-binding datasets with reference OCR set from this study

**TABLE S5: Gene-OCR associations**. Gene-OCR correlations from FigR (p<0.05) (Association = 2 in table) or other OCRs with 15kb of gene TSS (Association = 1 in table)

**TABLE S6: Topic Assignments**. OCR membership in each topic

**TABLE S7: Topic GREAT Gene Set Enrichment** Results related to Figure 3E. Gene Ontology gene sets (from GREAT ^76^) significantly enriched among regulatory regions in each topic. Values in table indicate fold change of enrichment relative to background.

**TABLE S8: IL2-dependent OCRs** Results related to Figure 5G. A) Fold Change (log_2)_ of aggregated accessibility in treated versus untreated cells, matched for cell state (rTreg or aTreg) for OCRs in Fig 5G. Each column is a different biological replicate. B) HOMER TF motif enrichment statistics in each IL2 response cluster (from Fig 5G). C) -log_10(_FDR) of Topic OCR enrichments in each IL2 response cluster (from Fig 5G).

**TABLE S9: Treg Regulatory Network** A) NonTSS Topic Motif Enrichment. B) TSS Topic Motif Enrichment. C) NonTSS Topic AME, unfiltered. D) NonTSS Topic AME, overlap with enrichment. FoxP3-dependent differential accessibility in topics for rTreg, all AME. F) FoxP3-dependent differential accessibility in topics for rTreg, enrichment-AME overlap. G) FoxP3-dependent differential accessibility in topics for aTreg, all AME. H) FoxP3-dependent differential accessibility in topics for aTreg, enrichment-AME overlap. I) FoxP3-independent differential accessibility in topics (KO vs Tconv), all AME. J) FoxP3-independent differential accessibility in topics (KO vs Tconv), enrichment-AME overlap. K) Motif Grouping Annotation Order. Note: for FoxP3-dependent changes, a value of 300 corresponds to grey cells in Fig 6 and S7, where there is a significant topic AME (FDR<0.10) but no FoxP3-dependent change (FDR<0.05).

**TABLE S10: Additional OCRs**. Information on additional OCRs used in F1, colon/spleen, and IL2 analyses.

