## Supplementary material for "An interwoven network of transcription factors, with divergent influences from FoxP3, underlies Treg diversity": Computational Notes

**Computational Note 1: Further Details about Ensemble Topic Modeling**

To learn coordinated programs of accessibility across Treg single cells, we used Latent Dirichlet Allocation ("topic modeling") with WarpLDA as implemented in cisTopic[1]. To arrive at a set of topics that would be robust across datasets and conditions, we sought to address two limitations in the default usage of cisTopic. First, as probabilistic models are inherently stochastic, the results vary from run to run. We addressed this variability by using ensemble modeling, learning several models with different initializations from the same data to construct a consensus set of reproducible topics. Second, while topic modeling learns continuous probability distributions for OCR membership in each topic, many downstream analyses rely on discrete peak sets. The original authors of cisTopic provide some guidance on how to binarize topics by setting a fixed threshold to select top OCRs after fitting to a Gamma distribution. However, each topic has slightly different probability densities, with some having most density concentrated in a few OCRs with others having probabilities more evenly distributed across many OCRs. Fixed thresholds do not incorporate these differences in topic OCR probability distributions. To address this limitation, we developed a heuristic approach that identifies OCRs specific to consensus topics, with flexibility to incorporate different topic assignment probability distributions. We provide additional details on our final approach below.

Ensemble modeling is a well-established approach for extracting robust signals from stochastic machine learning models[2,3]. To create a set of reproducible topics using ensemble modeling, we began by running 10 models with different initialization seeds on the Treg scATAC-seq dataset. For each model run, we chose the number of topics based on minimization of perplexity and maximization of the second derivative of model log likelihood curve, as recommended in the original cisTopic paper (Fig C1A). In cases where there were two solutions with approximately equivalent metrics for model fit, we opted for the model with the greater number of topics (with median of 16 topics per model), reasoning that redundant topics could later be collapsed when identifying consensus patterns (Fig C1A). Next, to identify reproducible patterns, we compared topic OCR probability distributions using Jensen-Shannon divergence, a symmetric measure for the similarity between probability distributions. Using these divergence values, we clustered topics from individual runs into consensus topics with Affinity Propagation Clustering (Fig C1B)[4,5]. Supporting their reproducibility, all resulting topic clusters had representation from a median of 8 separate individual model runs.

Next, we identified representative OCRs for each topic cluster, looking for OCRs that were coherently represented across all constituent topics per topic cluster but not in topics from other topic clusters. To do so, for each individual topic, we identified which OCR topic membership probabilities cumulatively contributed to the top 75% of the total membership probability. Formally, we looked for all OCRs, $x$, with probability $\geq$ a threshold $X$, such that $P(x \geq X) = 0.75$. By using this metric related to the cumulative distribution function, topics with different probability densities would have different numbers of selected OCRs. We calculated how frequently each OCR was in this top 75% for topics within each cluster, using non-cluster topics as the background comparators. OCRs with within-cluster topic frequencies that exceeded the $99^{th}$ percentile of non-cluster topic frequencies were chosen as the representative OCRs per topic. To identify topics distinguishing Treg from Tconv, we repeated this consensus topic selection procedure after running models on data with both cell types. We supplemented our Treg consensus topics developed above with two Tconv-specific topics (Topics 16 and 17) (Fig C1C). Our final solution had 17 topics, with $1.5\text{x}10^4$ to $3.9\text{x}10^4$ OCRs each (Fig C1D).

The final topic solution was robust and reproducible. Topics learned from running a new set of 10 model runs on the same Treg data could be readily assigned to a corresponding pattern in the consensus topics above (Fig C1E). Furthermore, the relative variance explained by each topic across two independent spleen datasets was highly reproducible (Fig 4B). Topic results were independent of the choice of threshold used to select representative OCRs per topic: comparisons between experimental conditions (Treg vs Tconv, FoxP3 KO vs WT, and Colon vs Spleen) were consistent across topic OCR selection thresholds (Fig C1F). Overall, our ensemble topic modeling approach enabled us to identify reproducible and robust Treg accessibility patterns that carried across datasets and experimental conditions.

**Computational Note 2: Topic-specific Allelic Motif Effect Calculations**

We employed a two-step strategy to identify causal contributions of motifs to each topic. First, we identified the “relevant” cells for each comparison, by looking for single cells with preferential accessibility of topic OCRs containing the candidate motif. Second, we aggregated B6- or Cast- allele-specific reads from only these relevant cells for input into the AME calculation to test the causal contribution of each motif to the topic of interest. The resulting outputs provided the final topic AME score.

To identify relevant cells for each comparison, we used chromVAR to calculate the per-cell, bias-corrected relative accessibility of topic OCRs containing each motif. We fit a two-component Gaussian mixture model (GMM) to these chromVAR scores to separate cells into a positive and negative set. To overcome noise in each model fit, we fit 10 separate GMMs for each comparison, retaining only those cells assigned to the positive set in all 10 model fits. Finally, we aggregated the corresponding B6- or Cast- allele-specific reads from the positive cell set for input into the AME calculation.

Once we had a set of aggregated reads corresponding to the relevant cells to be considered, we focused on OCRs from each topic of interest. We adapted motif analysis computations described in recent work[6,7]. Briefly, we created Cast-specific OCR sequences by modifying B6 OCR sequences with Cast-specific SNP and indel variants in vcf files from the Mouse Genome Project[8]. We then scanned B6 and Cast OCR sequences for all *Mus musculus* motifs from the cis-BP database[9] using FIMO[10] (retaining matches with $p < 10^{-4}$). To identify where variants affected motif matches, we looked for motif instances with different p values on B6 vs Cast alleles. For each OCR, we also calculated the $\log_2$ Fold Change in chromatin accessibility between B6- and Cast- specific aggregated reads (filtering out loci with fewer than a mean of 5 counts across the two alleles). For each motif, we calculated the AME score as the difference in $\log_2$ (B6/Cast) allelic bias in OCRs with stronger motif matches on the B6 allele vs on the Cast allele. In this framework, a positive score corresponds to a positive effect on chromatin accessibility and a negative score to a negative effect on chromatin accessibility. To assess significance of this difference, we used a two-sided t-test and applied the Benjamini-Hochberg false discovery rate procedure, retaining effects with FDR<0.10. We restricted analyses to motifs of TFs expressed in Tregs as defined from previous bulk and single cell RNA-seq datasets. The final, refined Treg TF network consisted of motifs with both significant enrichment (FDR<$1x10^{-10}$) and AME for a given topic (Fig 5B). All motifs with significant AME, regardless of corresponding enrichment, are also provided in Figure S5D.

### COMPUTATIONAL NOTE FIGURE LEGEND

**Figure C1: Details of Ensemble Topic Modeling.** A) Model perplexities for two examples of cisTopic model runs. Examples are shown for where model with greater number of topics is local (top) or global (bottom) minimum. Selected model indicated by red X. B) 1-(Jensen-Shannon Divergence) of topic OCR probability distributions for topics from 10 independent cisTopic model runs. Annotation colors indicate individual model runs (outside colors) or topic cluster assignment after Affinity Propagation clustering (inside colors). C) Relative accessibility (chromVAR scores) of OCRs from each topic across Treg and Tconv single cells, visualized on UMAP of Treg and Tconv scATAC-seq data from Fig S1. D) Number of OCRs selected as representative per topic. Color indicates TSS vs distal OCRs per topic. E) 1-(Jensen-Shannon Divergence) of topic OCR probability distributions for topics from a new set (rows) of 10 independent cisTopic model runs compared with previous set (columns) of 10 models in (B). Row annotations indicate model run for new set of 10 models. Column annotations model run and topic cluster assignment after Affinity Propagation clustering for models from (B). F) $\log_2$ Fold Changes in per-topic aggregated accessibility for each of the indicated comparisons across different thresholds for selecting representative OCRs.

Figure C1

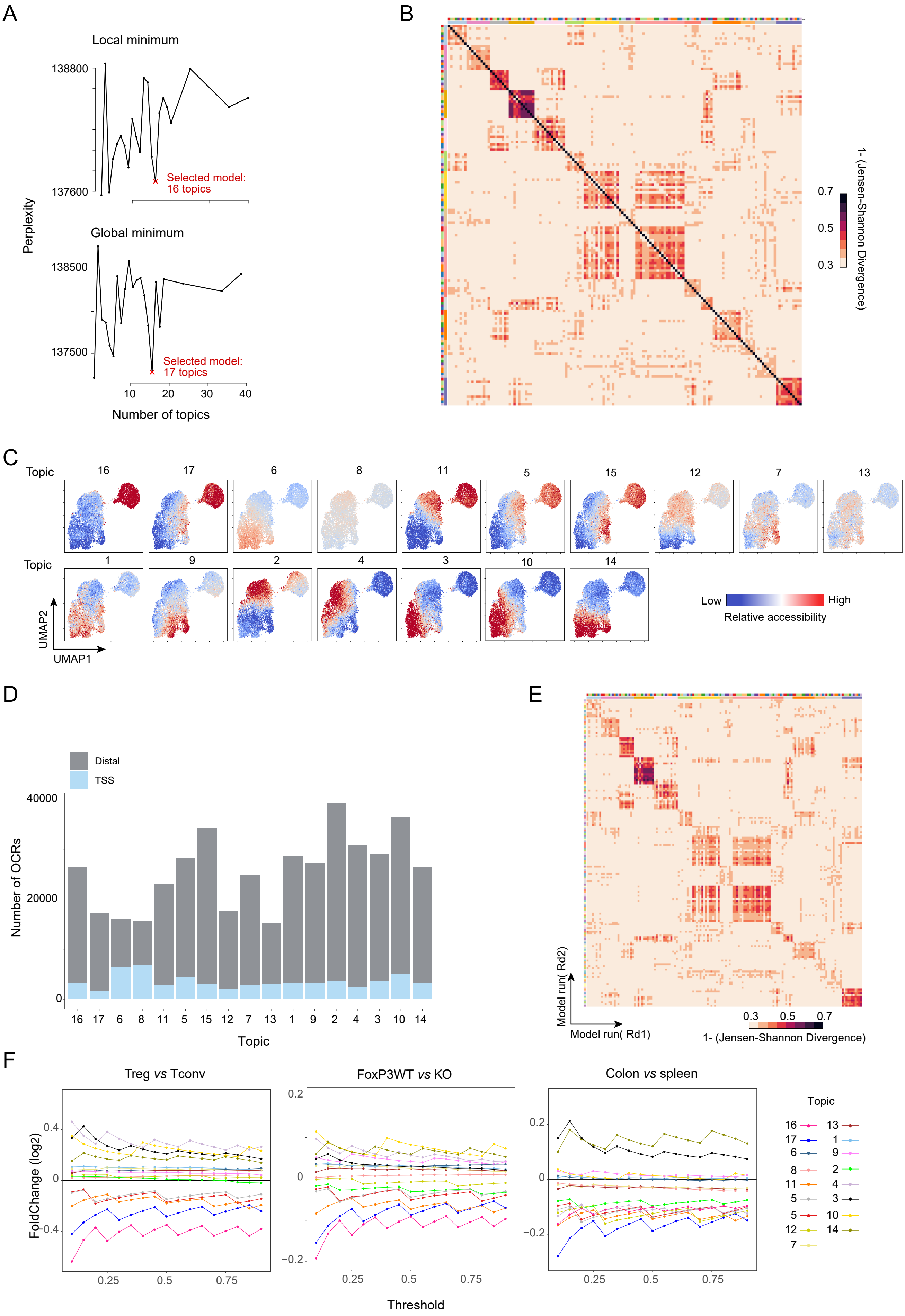
A
Local minimum
138800
137600
Selected model: 16 topics
Perplexity
Global minimum
138500
137500
Selected model: 17 topics
10
20
30
40
Number of topics
B
0.7
0.5
0.3
1- (Jensen-Shannon Divergence)
C
Topic
16
17
6
8
11
5
15
12
7
13
Topic
1
9
2
4
3
10
14
UMAP2
UMAP1
Low
High
Relative accessibility
D
Distal
TSS
40000
20000
0
Number of OCRs
16
17
6
8
11
5
15
12
7
13
1
9
2
4
3
10
14
Topic
E
Model run( Rd2)
Model run( Rd1)
0.3
0.5
0.7
1- (Jensen-Shannon Divergence)
F
Treg vs Tconv
FoxP3WT vs KO
Colon vs spleen
0.4
0
-0.4
0.2
0
-0.2
0.2
0
-0.2
FoldChange (log2)
0.25
0.5
0.75
Threshold
Topic
16
17
6
8
11
5
5
12
7
13
1
9
2
4
3
10
14
